# Contextualizing gene expression with feature rich graph neural networks

**DOI:** 10.1101/2025.03.17.643816

**Authors:** Steve S. Ho, Ryan E. Mills

## Abstract

Variation in gene expression arises from the interplay between chromatin architecture, epigenetic marks, transcription factor binding, and regulatory elements – none alone capture the full complexity of the regulatory landscape. We introduce Omics Graph Learning (OGL), a deep learning paradigm that integrates multi-omics data into a network of biological entities. Leveraging over 950 curated datasets, OGL achieves highly accurate gene expression prediction across 20 cell lines and tissues. Through > 81 million *in-silico* perturbations, we uncover tissue-specific patterns of feature utilization, reveal non-additive interactions among epigenetic marks, and identify regulatory elements with disproportionate influence on gene expression. Our findings highlight complex synergistic and antagonistic relationships among molecular features that vary by cellular context. By bridging predictive accuracy with biological interpretation, OGL provides a framework for deciphering multifaceted mechanisms that shape the transcriptome.

## Introduction

DNA sequence composition is an integral part of but not solely deterministic of phenotype – genes ultimately need to be expressed to carry out their function^1,2^. Gene expression emerges from a complex orchestration of molecular features such as chromatin architecture^3^, transcription factor binding^4^, regulatory element coordination^5^, epigenetic activity^6^, methylation^7^, and other interactors like non-coding RNAs^323^ and RNA-binding proteins^9^. Despite substantial progress in characterizing individual components of the regulatory landscape, we lack comprehensive frameworks that integrate these diverse elements to explain how they collectively shape gene expression patterns.

Deep neural networks have transformed genomic analysis by offering powerful tools to decode complexity through the automatic extraction of hierarchical features from raw data^10^. In genomics, neural networks are commonly used in sequence-to-function models that predict molecular assays from DNA sequence alone, effectively learning the regulatory grammar of the genome without explicit programming of biological rules^11–13^. To interpret these models, researchers use specialized techniques including gradient-based saliency^14^, explanations^15^, and *in-silico* saturation mutagenesis, which systematically perturbs inputs to reveal which nucleotides most significantly impact model predictions.

While convolutional neural networks^11,13,16,17^ and transformers^18,19^ have dominated this landscape, their inherent architecture constrains their ability to integrate distal regulatory information. Most predictive attributions from these models concentrate near promoter regions likely due to the significant imbalance between actual and candidate regulatory elements as genomic distance increases^20^. This architectural limitation mirrors a fundamental challenge in understanding gene regulation: many crucial interactions occur between elements separated by large linear genomic distance yet are brought into physical proximity through three-dimensional chromatin organization^21^.

Graph neural networks (GNNs)^22^ can directly address this limitation by explicitly encoding relationships between genomic elements. Whereas sequence-based models rely on a spatial inductive bias, GNNs can represent the interactions that characterize gene regulatory networks as direct relationships. Previous genomic GNN implementations segment the genome into fixed-size bins which are connected via chromatin interaction data^23–25^. However, this binning approach introduces distortions: gene bodies get fragmented across multiple bins and multiple distinct regulatory elements get compressed into single bins, hindering interpretability.

We introduce Omics Graph Learning (OGL), a modeling paradigm that represents the genome as a network of biologically meaningful entities where each node corresponds to a complete genomic element, such as a gene, enhancer, or promoter, providing a discretized framework. Our models achieve highly accurate expression prediction across diverse cell lines and tissues, serving as a foundation for deeper regulatory insights. OGL’s interpretability-first design decouples molecular features from sequence effects, enabling systematic interrogation across multiple scales, with broad-scale analyses like gradient saliency and feature ablation, and fine-scale investigations through individual node perturbations. Over 81 million *in-silico* perturbations reveal a regulatory landscape where most elements confer modest effects while a small fraction (∼0.05%) exert disproportionate influence. OGL uncovers tissue-specific regulatory patterns, non-additive relationships between epigenetic marks, and the context-dependent ability of elements to function as both activators and repressors. By bridging the gap between prediction and interpretation, OGL challenges reductionist paradigms in gene regulation, offering insights into how molecular features emerge as expression.

## Results

### Feature-rich graph deep neural network model

This work operates on the premise that a performant neural network will approximate the most pertinent biological interactions to accurately predict expression. Thus, our experimental design combines two critical elements: a highly accurate expression prediction model to ensure we capture valid biological signal with an interpretability-first approach to enables systematic *in-silico* experimentation to dissect the molecular determinants of gene regulation. To achieve this, we utilize GNNs to predict gene expression by combining regulatory element annotations, 3D chromatin assays, and a rich set of omics and genomic features. OGL models the relational inductive bias between genomic elements by explicitly encoding biological data in the graph structure, allowing the GNN to learn complex non-linear interactions between these elements.

OGL produces sample-specific, sequence-agnostic graphs derived from multi-omics data (Extended Data Fig. 1). Each graph is a network of genes and regulatory elements connected through chromatin contacts. To ensure high-confidence regulatory elements, we use an intersection of the ENCODE SCREEN v3 registry and EpiMap candidate *cis*-regulatory elements (CREs). We ensemble chromatin data from orthogonal methods to generate comprehensive, sample-specific catalogues of 3D contacts. The graph nodes (genes and regulatory elements) are connected by these contact catalogues, and we enrich expressivity by connecting each node to all others within a context window to effectively capture each element’s local genomic neighborhood (see Supplementary Information). Nodes are augmented with comprehensive feature vectors incorporating both tissue-specific genome tracks (such as histone modifications) and static genomic features (such as GC content and reference repeat overlap) (Supplementary Table 1).

Node feature vector are concatenated with a binned positional encoding which informs the model of relative positioning for each feature. The graph is fed into GNN layers utilizing a dynamic attention mechanism to attend each node with all of its neighbors (GATv2)^26^. Representations are non-linearly transformed with the GELU^27^ activation function and gradient flow is augmented with a distinct-source residual connection. Fully connected layers propagate the learned signals to separate task heads which simultaneously regress expression values (log_2_ TPM) and perform classification (high or low gene expression). We note that this only describes the most performant architecture of the study; OGL itself implements flexible graph construction and couples it with a modular GNN architecture. Supplementary Table 2 details the final model architecture used for this study and the flexible architecture and graph construction are further detailed in Extended Data Fig. 2 and the Supplementary Information.

**Fig 1.**
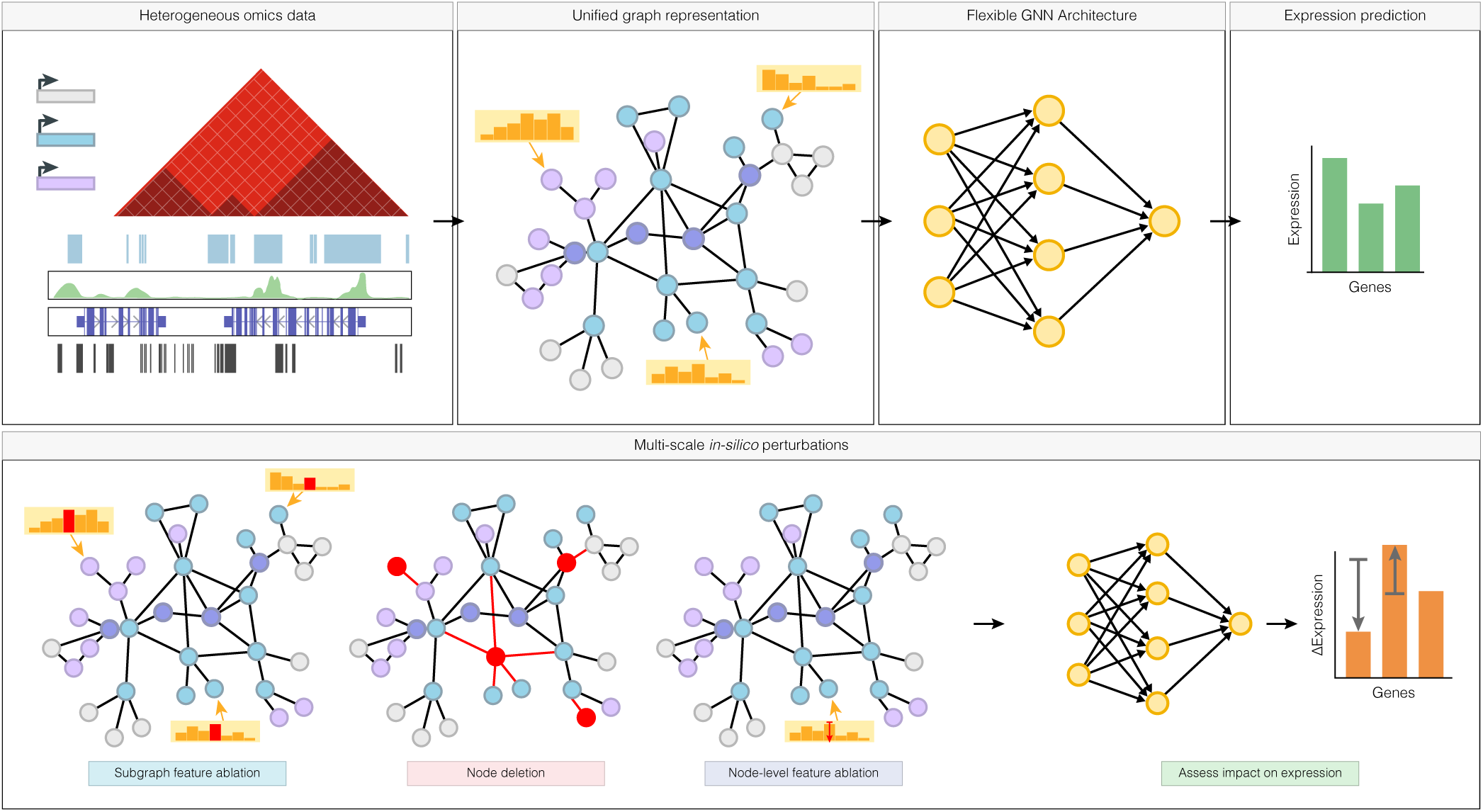
Overview of OGL pipeline. OGL constructs graphs from rich feature data to generate a unified representation across modalities. The graphs are used as inputs into sample-specific GNN models that regress gene-level expression levels. OGL’s design facilitates interpretable multi-scale perturbation experiments at the subgraph and node levels, enabling systematic perturbation and ablation studies to query the impact of genomic elements and molecular features on expression.

### OGL accurately predicts sample-specific gene expression

We first developed models for K562 due to its well-characterized regulatory landscape and high-resolution 3D chromatin data. OGL accurately predicts gene-level expression (Pearson’s *R* = 0.896) on hold-out test genes (Fig. 2a). Although performance is difficult to compare directly due to varying data processing and model design, OGL produces state-of-the-art performance compared to other deep learning models predicting gene-level expression in K562 (Supplementary Table 3). We stress that OGL was designed as an interpretability-first framework and that its strong predictive performance emerges as a byproduct of biologically meaningful design choices rather than purely task-oriented optimization.

**Fig 2.**
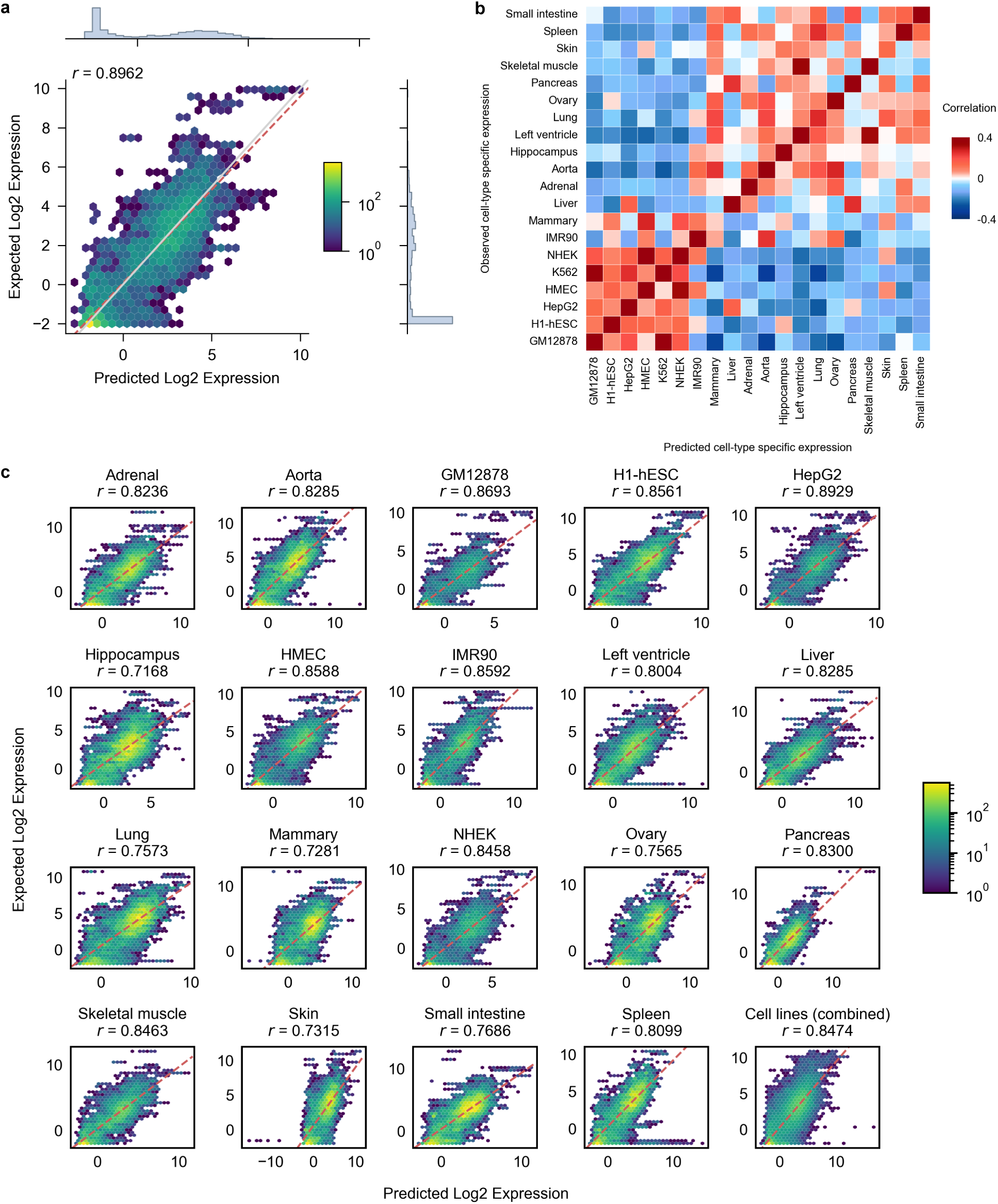
OGL accurately predicts expression across cell-lines and tissues. a, OGL produces SOTA performance in predicting gene-level RNA-seq within K562. Test set performance is measured as Pearson correlation of expected versus predicted gene expression (log2(x + 0.25)) on hold-out genes (chromosomes 8 and 9). A best fit line (gray) and a regression line (red) are displayed for comparison. b, Heatmap of expression on holdout genes, shown as correlations between sample-specific expression profile as measured by log_2_(fold change) value over the sample-type average and the predicted log_2_(fold change). c, Gene-expression predictions on holdout genes are highly correlated with expected log2(TPM) values in across different cell lines and tissues (average correlation across all models = 0.83).

We extended OGL to a total of 20 cell lines and tissues. OGL captures sample-specific expression patterns, showing higher similarity between biologically related cell types and tissues on hold-out test genes (Fig. 2b) and accurately predicts expression across all samples (Fig. 2c). The complete set of 20 models achieves an average Pearson’s *R* of 0.83, while the cell line models attain an average *R* ∼0.87: this performance gap likely results from differences in underlying data quality. We observe moderate correlation (Spearman’s ρ = 0.695; Supplementary Fig. 1) between the total amount of chromatin contacts used for graph construction and performance. However, this correlation understates the quality disparity, as tissue-specific contact data typically exhibits lower resolution and technical quality compared to cell line data, which are limitations not adequately captured in processed contact counts alone. These results underscore the critical importance of high-quality, comprehensive input data for achieving optimal model performance.

The flexibility of OGL allowed us to explore a combined model incorporating graphs from all cell lines (Fig. 2c). While this generalized model performed well (Pearson’s *R* = 0.847) it did not exceed performance of the individual models. We focused on individual models for all subsequent analyses to maximize the probability of extracting meaningful biological insights.

### OGL focuses on pertinent sample-specific features for prediction

To better understand how OGL leverages molecular features for expression prediction we computed raw gradient-based saliency scores for all possible nodes in the graph which reflect the sensitivity of the model predictions with respect to each feature, providing a relative measure of feature importance (Supplementary Information). Per-feature averaged gradient magnitudes revealed distint sample-specific patterns (Extended Data Fig. 3a). Notably, comparing our most performant models (K562 and HepG2) reveals marked differences in which features show high sensitivity, suggesting that OGL effectively prioritizes sample-specific features. Global saliency maps (Extended Data Fig. 3b) highlight model-specific differences: the K562 model exhibits narrower, focal peaks of gradient sensitivity, whereas the HepG2 model shows a more distributed pattern of sensitivity across the feature landscape.

We stratified the saliency maps by node class (Supplementary Fig. 2). As expected from previous work^20,28^, we fine genes features exhibit highest gradient magnitudes, followed by promoters, then enhancers. Inspecting high-sensitivity nodes across models suggests a higher degree of tissue specifity in enhancers than promoters which aligns with previous work^28^ (Supplementary Fig. 3). Although many nodes show modest gradients, we identified subsets with strongly elevated sensitivity, especially among genes. Notably, while our models only consider protein-coding genes during loss calculation, we found substantial gradients from various non-coding RNA classes (Supplementary Fig. 4), including antisense transcripts, long intergenic non-coding RNAs, and pseudogenes, all of which have known roles in gene regulation^29–31^. This suggests that OGL models effectively integrate information from broader genomic interactions rather than solely relying on features from protein-coding genes used for prediction.

We then investigated whether OGL models identify distal enhancers that influence gene expression. We analyzed gold-standard CRISPR interference (CRISPRi) element-gene pairs from Gschwind et al.^32^ focusing on substantial effect pairs (≥ 10% expression change) (Supplementary Information). Correlating gradient saliency with CRISPRi effect sizes revealed a weak positive correlation with statistically insignificant pairs, (Supplementary Fig. 5a, Spearman’s ρ = 0.188) and a pronounced negative correlation with significant pairs (Supplementary Fig. 5b, Spearman’s ρ = −0.304). This negative correlation indicates that our model assigns higher feature gradients to elements with stronger repressive effects, suggesting OGL effectively prioritizes functionally relevant regulatory relationships. Further analysis revealed minimal correlation between individual feature scores and predicted CRISPRi effects (mean Spearman’s ρ = 0.033), indicating that OGL evaluates the collective feature importance rather than relying on any single dimension to prioritize putative cCREs.

To gauge contributions, we retrained K562 models by randomizing individual molecular features while keeping others static (Supplementary Fig. 6). The baseline model with all features preserved performed best (average Pearson’s *R* = 0.893 over three seeds) with the most stable performance. Across randomized models, no single feature randomization significantly degraded model performance, indicating that each feature contributes incremental signal to the predictive capacity. These results suggest that the full feature combination is necessary to capture the complete regulatory landscape. When all features were simultaneously randomized, performance dropped substantially (Pearson’s *R* = 0.008), confirming the model captures biological information rather than overfitting or exploiting spurious correlations.

### Broad-scale perturbations reveal non-additive, directional effects of molecular features

Gradient saliency reflects sensitivity but does not inform about the direction of influence, importance of feature magnitude, or the relationship between feature and output beyond sensitivity. We turned to broad-scale *in-silico* perturbations to gain deeper insight into mechanisms OGL learns, performing ablation studies where we completely removed molecular features (e.g. removing all CpG methylation) from each gene’s *K*-hop subgraph to measure the impact on expression prediction (Fig. 3a). In total, we performed > 13 million ablations at the subgraph scale.

**Fig 3.**
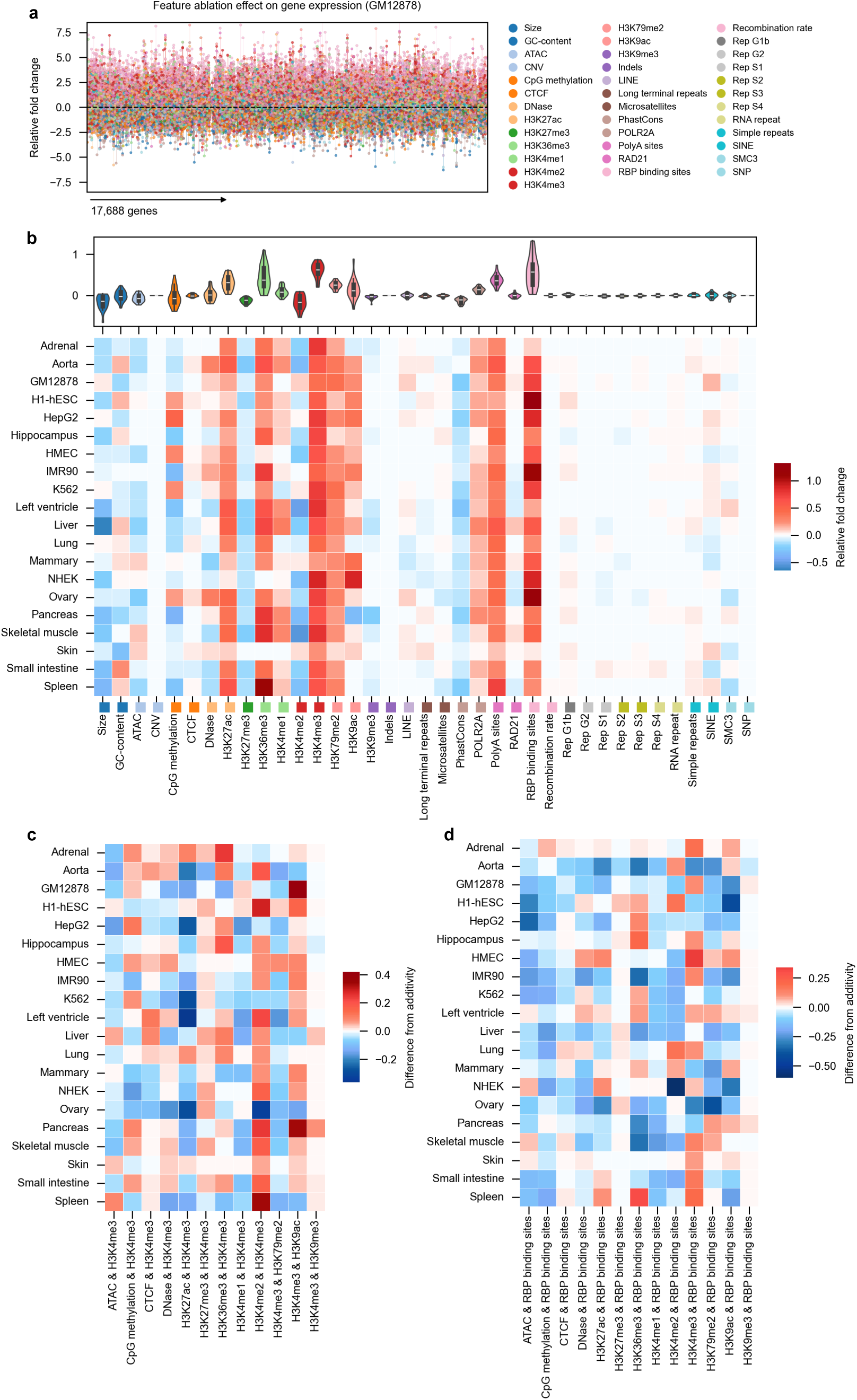
Embeddings encode known biological relationships. a, The semantics embedded into vectors recapitulate broad scientific relationships through by vector operations that look for similarities in Euclidian space. The similar direction and magnitude of these vectors suggests that the embeddings capture meaningful relationships. For example, as “anaphase” is characterized by “separation”, “metaphase” is characterized by “alignment”. The depicted relations are of broad biological analogies, b, known gene-gene interactions, and c, gene-disease associations. d, Co-occurrence predictions between 10,000 randomly sampled gene-gene interactions that overlap GO pairs and. e, gene co-essentiality.

The feature ablation map contrasts strongly with the saliency map (Fig. 3b): while saliency scores showed sample-specific patterns, the ablation map revealed remarkable consistency across samples. Sample-specific features like histone modifications, CpG methylation, and open chromatin regions generally had larger predicted fold changes compared to genome-static tracks like recombination rate or evolutionary conservation scores, with exceptions. Notably, removal of RNA-binding protein binding site clusters had the largest impact across samples, complementing previous work that integrates post-transcription annotations for expression prediction^33^. These results demonstrate the importance of integrating both genome-static and sample-specific molecular features when modelling gene expression.

We quantified the impact of genome-static features by identifying genes that showed significant expression changes (|FC| > 0.1) upon feature ablation despite having no direct overlap of that feature within the gene body itself. Microsatellite annotations had the most substantial impact, affecting 45,000 genes (15.73% of genes across all models) exclusively due to their removal in regulatory neighborhoods, consistent with emerging evidence of their role in transcription^34^. Other features demonstrating regulatory influence included long terminal repeats (7.92%) and indels (5.77%). These results illustrate that OGL effectively captures how genomic features within the broader regulatory landscape can influence gene expression indirectly.

We observed unexpected directional effects upon ablating certain node features. For instance, removal of ATAC open chromatin regions generally led to decreased expression, consistent with open chromatin facilitating transcription, yet ablation of H3K4me3 regions produced modest increases in expression (Fig. 3b). To investigate further, we performed joint feature ablations focusing on combinations with H3K4me3 (Fig. 3c) and RBP binding site clusters (Fig. 3d). Our analysis revealed sustained deviations from additivity for multiple feature combinations. Notably, the pronounced non-additivity for H3K4me2 and H3K4me3 illustrates their cooperative relationship^35,36^ (Supplementary Fig. 7). Consistent sub-additive effects emerged from combining histone ablations, exhibiting distinct tissue-specific patterns (Fig. 3c). Similarly, feature combinations involving RBP binding site clusters displayed non-additive behavior with varying degrees of deviation and tissue specificity (Fig. 3d). This suggests that the model attributes nuanced function to histone modification signals, and aligns with evidence that regulatory elements possess both activating and repressive potentials, even those marked as “active”^37–41^. These findings demonstrate that our models capture how molecular features influence gene expression synergistically rather than independently.

To assess the biological significance of these perturbations, we performed gene set enrichment analysis on the top 100 genes most affected by each ablation. The results largely met expectations: H3K27me3 ablation enriched for DNA binding processes^42,43^ (Supplementary Fig. 8), while ablations in IMR90, skeletal muscle, and H1-hESC enriched for processes involved in extracellular matrix organization, motor unit control, and pluripotency and development, respectively (Supplementary Fig. 9).

### OGL enables fine-scale perturbation of expression models

Next, we assessed the regulatory impact of individual nodes by systematically deleting each node in the *K*-hop subgraph of accurately predicted genes (see Methods) and measuring the change in predicted gene expression, producing a comprehensive map of > 51 million perturbations. The global perturbation landscape (Fig. 4) reveals distinct patterns across different node classes. Gene perturbations (Fig. 4a) exhibit the most substantial effects with an average absolute relative fold change (|FC|) of 0.0335, followed by promoters (Fig. 4b; average |FC| = 0.0071), while enhancers (Fig. 4c; average|FC| = 0.0026) exhibited lower effects. This pattern is more pronounced when filtering for strong effects (|FC| > 0.01), with genes showing an average effect of 0.0748 followed by promoters (0.0415), and similar effects for dyadic elements (0.0290) and enhancers (0.0289). Although the visualization suggests greater variability in enhancer effects, this is due to their larger number in the dataset, resulting in a denser distribution. Overall, only about 0.2% of perturbations (107,988 nodes) induced changes of ≥10%, and 0.02% (11,441 nodes) resulted in changes of ≥50%. These findings suggest that OGL learns gene regulation emerges from the cumulative influence of many elements, while a few influential regulators drive more substantial effects^44^.

**Fig 4.**
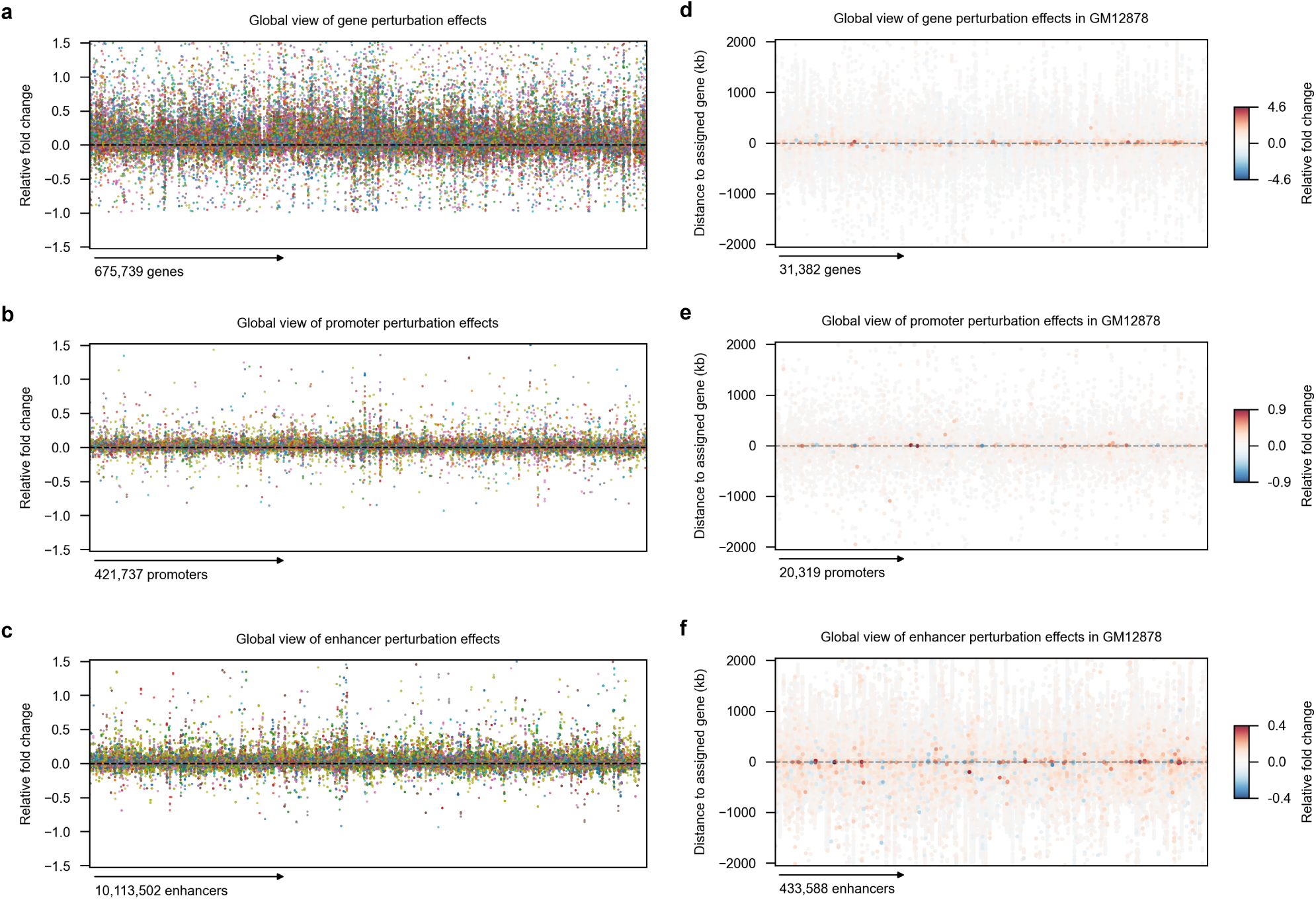
A map of *in-silico* perturbation experiments. a, Each point represents the predicted relative fold change in predicted gene expression upon deletion of a connected node. Values were computed as the max effect of the specified node deletion across all possible subgraphs for which that node is present. Colors indicate the different sample that each perturbation derives from. Elements are ordered by chromosome location (1-22) from left to right. Node types perturbed in each panel a, genes. b, promoters, c, enhancers. d, Global view of gene perturbation effects in GM12878, showing the relationship between genomic distance (y-axis, kb) to the assigned gene and relative fold change in expression (color intensity) for 31,382 gene nodes. Distances are shown both upstream (negative values) and downstream (positive values) of the gene. e, Spatial distribution of 20,319 promoter perturbation effects in GM12878, demonstrating more distributed regulatory influence. f, Distribution of 433,588 enhancer perturbation effects in GM12878, showing the broadest spatial reach across the regulatory landscape. Point intensity corresponds to the magnitude of relative fold change upon node deletion, with red indicating increased expression and blue indicating decreased expression.

We investigated sample specificity of the perturbations by calculating tissue-specific scores (Tau) for each node based on their perturbation effects across samples. Perturbed nodes skew strongly toward higher Tau scores (Supplementary Fig. 10). Notably, elements with Tau values above 0.8 exhibited higher perturbation effects than those with lower specificity scores.

We examined the spatial distribution of effects relative to target genes. Influential gene nodes were commonly in close proximity to assigned genes (Fig. 4d), influential promoters exhibited a slightly broader spatial pattern (Fig. 4e), and influential enhancers displayed the most extensive spatial distribution (Fig. 4c). The progression from highly localized gene effects to the more distributed influence of promoters and the broad spatial patterns of enhancers illustrates how OGL models capture the hierarchical organization of the regulatory landscape.

The distribution of perturbation effects was asymmetric with positive and negative fold changes observed across all node classes (Fig. 4). This asymmetry mirrors findings from CRISPRi screens where a substantial proportion (41%) of enhancer perturbations result in increased expression^32^. We sought to see if our model was capturing putative silencers^45^ and binarily classified elements as either repressive or activating based on the direction of perturbation effect. Among the 403 experimentally verified silencers in our K562 dataset, OGL successfully recalled 254 (63%; binomial test *p* < 1e^−6^). This suggests that our approach effectively balances the activating and repressive effects across numerous interacting elements and aligns with growing evidence of context-dependent regulatory activity that can be both activating or repressive^37–40,45–50^

### Node-level ablations reveal context-specific regulatory complexity

Building on our node perturbation analyses, we performed a more granular investigation by systematically removing individual features at the node level. For the 10,000 elements with the largest predicted perturbation effect in each sample, we selectively ablated individual features to measure their contribution to expression prediction (see Methods), performing > 7 million perturbations. Extended Data Fig. 4 illustrates the granularity of fine-scale perturbations, revealing signatures of feature utilization vary across the elements connected to a gene.

We averaged the node-level feature perturbation effects (Fig. 5a). Strikingly, node-level ablation of H3K27ac led to a decrease in expression, while ablation of H3K27me3 resulted in an increase— both effects that were obscured in our broad-scale analysis (Fig. 3b), underscoring potential bifunctionality. The distribution of feature-specific effects is markedly asymmetric across models: few features (H3K4me2 and conservation scores) exhibit consistent patterns, with most displaying element-level specificity. Fig. 5b compares predicted effects between GM12878 lymphoblastoid cells and H1-hESC embryonic stem cells revealing differential impacts of histone modifications and repetitive elements. Notably, removal of H3K4me3 at individual elements results in more pronounced expression decrease in H1-hESC cells, consistent with this mark’s established role in maintaining pluripotency^51^. These observations demonstrate that regulatory elements operate within complex, context-dependent networks rather than serving universal roles.

**Fig 5.**
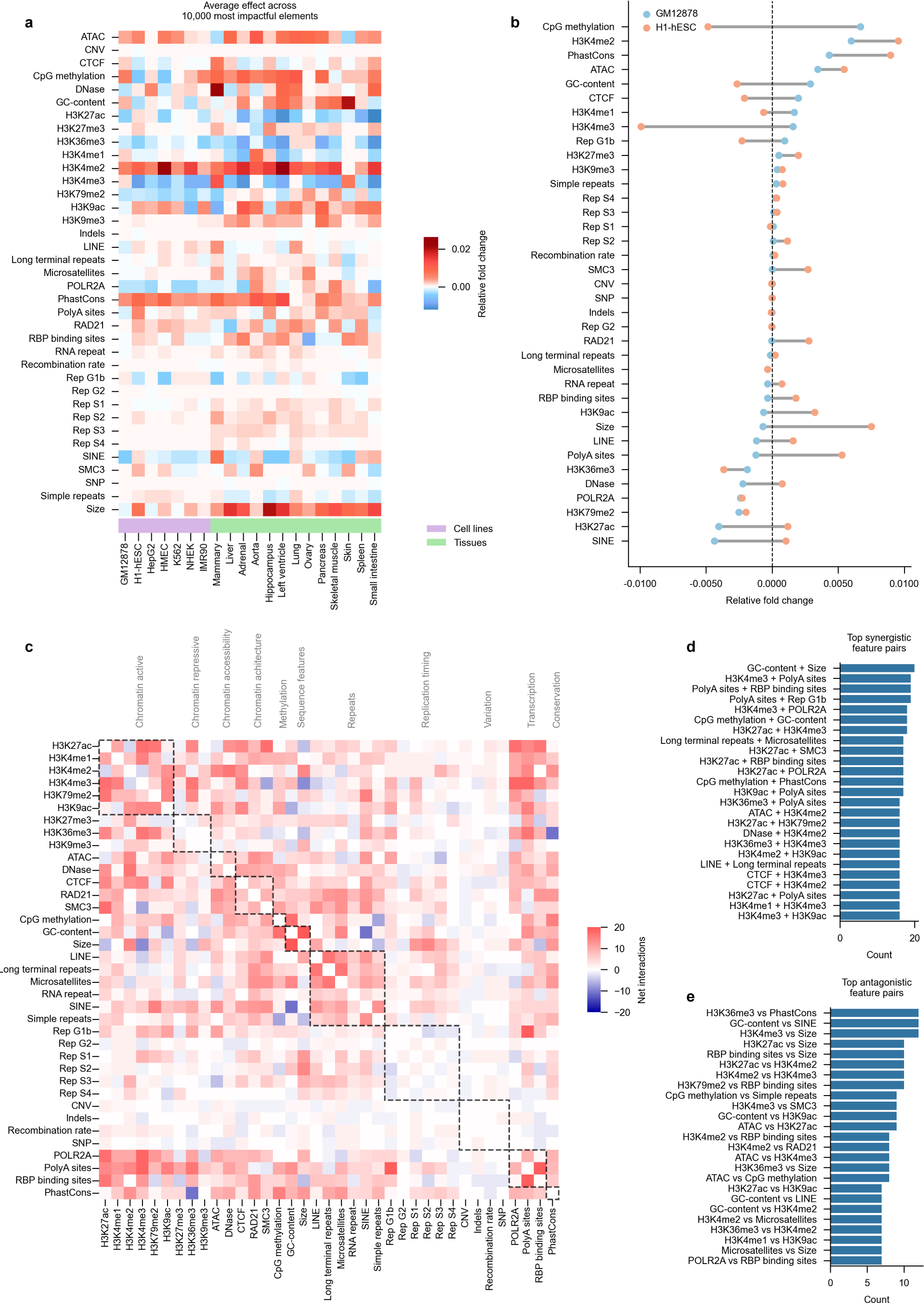
Feature interactions reveal cooperative and opposing regulatory relationships. a, Heatmap showing average effect per feature across the 10,000 most impactful elements in each tissue and cell line. Each row represents a sample, each column a molecular feature, and color intensity indicates the averaged relative fold change upon feature ablation. b, Dumbbell plot comparing the relative impact of node-level feature ablation between H1-hESC and GM12878 showing context-specific opposing effects. c, Interaction frequency matrix across all samples derived from partial correlations with functional categories (outlined by dashed lines). Red indicates net positive interactions (features with positive partial correlation), while blue indicates net negative interactions (features with negative partial correlation). d, Most common synergistic feature pairs identified across all samples, where both features influence expression in the same direction. e, Most common antagonistic feature pairs, which influence expression in opposite directions.

We then investigated interaction patterns among features to determine how different molecular components cooperate or oppose each other in regulating gene expression. For each element-feature perturbation, we computed correlations for all feature pairs conditioned against all other features. Comparing feature-correlation patterns across cell types (Supplementary Fig. 11) reveals context-specific regulatory interactions, with cell types differing in both interaction directionality and co-regulation frequency. Next, we quantified interaction frequencies by summing synergistic (positively correlated) and antagonistic (negatively correlated) feature pairs across samples (Fig. 5c). Notably, while expected correlations emerge within functional groups (e.g., active chromatin marks), many interactions span different groups, indicating that regulatory relationships frequently transcend conventional boundaries. For instance, H3K27ac emerges as a top synergistic and antagonistic partner with H3K4me2 and H3K4me3 (Fig. 5d,e), which are both active marks, highlighting functional duality and context dependency of these interactions. Together, these results illustrate that OGL can illuminate subtle, context-specific regulatory relationships by enabling *in-silico* perturbation of diverse genomic features, offering a comprehensive view of the expression landscape.

### OGL enables integration with sequence-to-function models

A significant advantage of OGL’s interpretability-first approach is its ability to complement sequence-to-function models through feature decoupling. Joint-task models like Enformer predict multiple genomic tracks simultaneously from sequence, creating implicit couplings between features and expression outcomes, presenting a fundamental limitation: when a sequence variant affects both a regulatory mark and gene expression, it remains unclear whether the expression change results interactions involving the mark alteration.

OGL addresses this limitation by operating downstream of sequence and representing molecular features as discrete node attributes. This design enables a two-step interpretability process that creates precise mechanistic pathways (Fig. 6). First, sequence-to-function models identify variants that alter specific molecular features at regulatory elements. Then, OGL quantifies how those altered features affect gene expression through targeted ablations. Figure 6 demonstrates this integration with two examples. In K562 cells, *in-silico* mutagenesis (ISM) with Enformer identifies variants likely to impact CTCF binding at a specific enhancer (Fig. 6a). Subsequent ablation of CTCF at this enhancer within OGL predicts increases expression of the target gene *PMVK*. Similarly, in GM12878 cells, Enformer identifies variants affecting H3K4me1 at an enhancer, and OGL reveals that removing this mark decreases expression of *RXRA* (Fig. 6b):

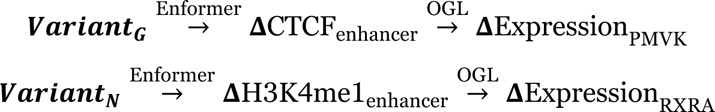

**Fig. 6.**
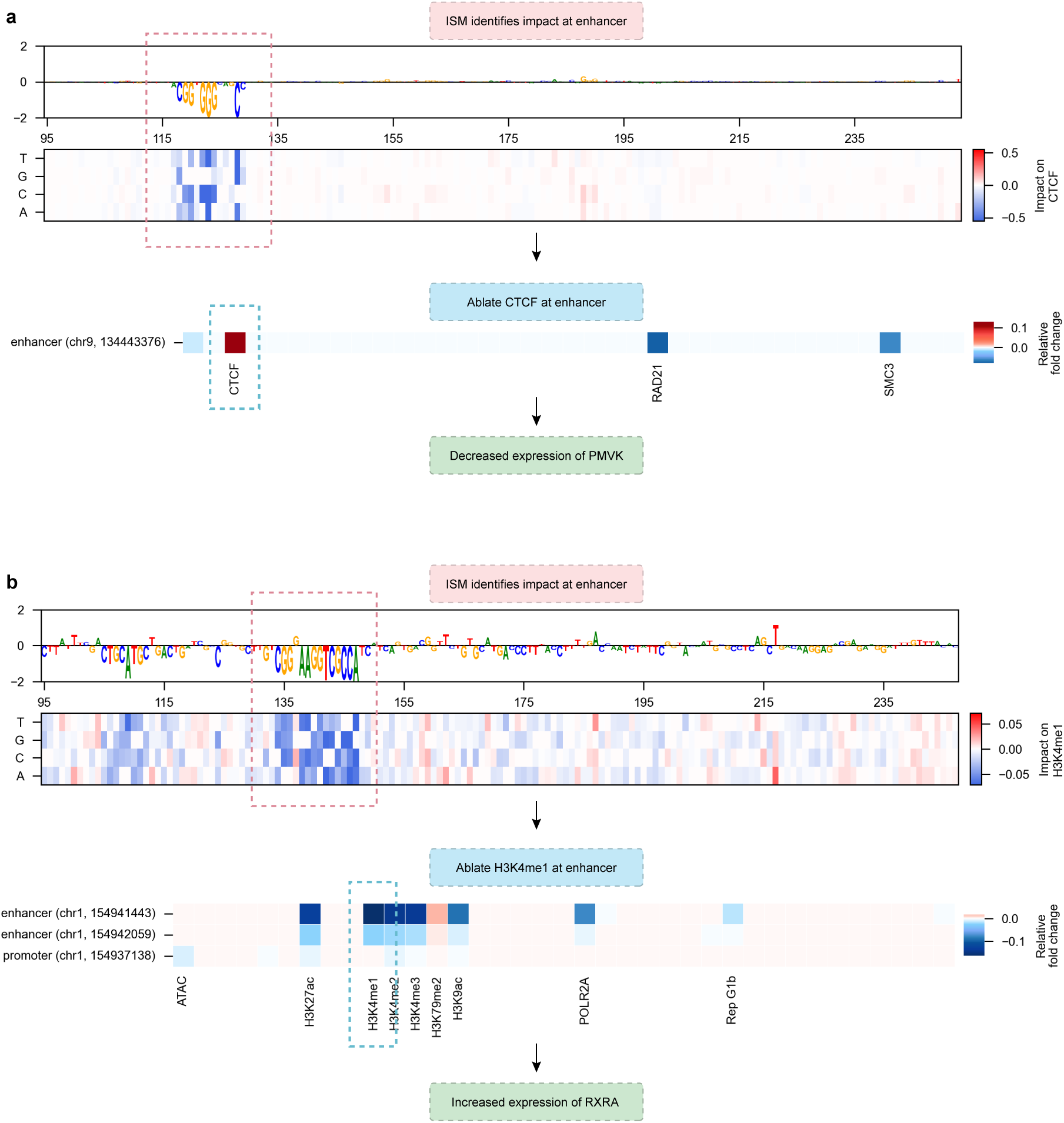
OGL enables integration with sequence-to-function models to predict impact from variant to expression. a, ISM with Enformer identifies variants most likely to impact CTCF at an enhancer in K562. OGL predicts removal of that CTCF at the enhancer increases expression of the target gene *PMVK*. b, ISM with Enformer identifies variants mostly likely to impact H3K4me1 at an enhancer in GM12878. OGL predicts removal of H3K4me1 at that enhancer decreases expression of the target gene *RXRA*.

These examples illustrate how the combined approach creates interpretable pathways from sequence variation, to molecular feature changes, to expression outcomes.

## Discussion

Omics Graph Learning (OGL) represents a departure from conventional sequence-based deep learning models by explicitly modeling genomic elements as networks of discrete biological entities rather than sequence bins. This approach allows us to interrogate gene regulation with granularity while achieving accurate expression prediction across diverse cell types and tissues. Our multi-scale interpretability framework reveals that OGL captures sample-specific regulatory patterns, with different cellular contexts prioritizing distinct molecular features. Feature ablation studies demonstrate complex, non-additive relationships among these features, underscoring the intricate synergy that shapes gene expression. From > 81 million *in-silico* perturbations, we observed that most regulatory elements confer modest effects on expression, while a small fraction (∼0.05%) exert disproportionately large impacts, mirroring observations from CRISPRi screens. OGL also captures both activating and repressive roles across regulatory elements, in line with growing evidence of context-dependent regulatory function^39,40,45,52^.

A key strength of OGL’s design is in its facilitation of interpretability methods. In practice, different attribution methods often yield conflicting signals^53,54^. When systematically comparing attention scores, perturbation effects, and saliency maps across our 20 models, we observed minimal correlation (Extended Data Fig. 5) between techniques. This discordance highlights a crucial pitfall: no single method captures the full picture of how a deep learning model makes predictions^53,54^. OGL’s systematic perturbation approach directly measures feature impacts without relying on assumptions from any particular attribution method.

Several key limitations remain. This work represents a curation-heavy specialized modelling approach opposed to parameter-maximized generalizability. Thus, OGL models rely heavily on the underlying quality of their input data, creating an unavoidable performance bias favoring well-characterized cell lines over tissue samples that lack extensive functional profiling and quality standards. Second, OGL utilizes static representations of molecular features which fail to capture the inherently dynamic nature of chromatin organization, histone modifications, and gene expression, oversimplifying the continuous, context-dependent states of these features. Additionally, the modeling framework imposes an implicit directionality that predominantly assumes histone marks are cause but not consequences of transcription^6^. Future models may benefit from bidirectional modeling that acknowledges the complex interplay between these states. Third, while we implement a multi-scale interpretability framework, the inherent complexity of neural network approaches necessitates rigorous experimental validation. The granularity of our perturbations currently exceeds the resolution of available experimental techniques, creating a validation gap where computational predictions outpace our ability to experimentally verify them at the same level of precision.

We recognize that the data-intensive nature of OGL invites significant implementation challenges. While we release model weights and the complete global perturbation map to maximize accessibility, we believe OGL’s primary impact should be conceptual rather than as a widely deployed tool. We demonstrate the value of integrative frameworks and highlight how conventional approaches that examine features in isolation might miss critical complexity. Our work challenges the reductionist paradigms that currently dominate the field – approaches that often characterize regulatory elements with binary descriptions and oversimplified annotations. Our hope is that shifting the field toward frameworks that embrace complexity rather than attempting to reduce the intricate regulatory landscape may ultimately prove more valuable than the specifics of our implementation.

In summary, OGL bridges predictive accuracy and biological insight by enabling systematic interrogation of regulatory mechanisms, enhancing our understanding of the complexities that shape the transcriptome.

## Funding

S.S.H. was supported by the NSF GRFP and the Michigan Predoctoral Training Program in Genetics (T32 GM007544).

## Competing interests

The authors declare no competing interests.

## Author contributions

S.S.H. conceived the project, curated the data, designed the pipelines, trained the models, performed all analyses, and wrote the manuscript. R.E.M. supervised the project.

## Data and materials availability

All code is available at https://github.com/sciencesteveho/ogl.

## Methods

### Data collection

We downloaded over 900 disparate tracks spanning both sample-specific and genome-static features, focusing on samples with data across multiple modalities. Our models span 7 cell lines (K562, IMR90, GM12878, HepG2, H1-hESC, HMEC, and NHEK) and 13 tissues (hippocampus, lung, pancreas, skeletal muscle, small intestine, liver, aorta, skin, left ventricle, mammary, spleen, ovary, and adrenal). Dataset accessions and download URLs are noted in Supplementary Table 4.

### Gene annotations

We used the GENCODE^55^ v26 reference for the hg38 reference genome and filtered for protein coding genes on chromosomes 1-22. We removed microRNA annotated genes unless the microRNA network is added during graph construction.

### Regulatory catalogue assemblage

We downloaded regulatory enhancer-like, promoter-like, and CTCF-only elements from the ENCODE SCREEN v3 registry^56^, and downloaded enhancer, promoter, and dyadic annotations from EpiMap^57^ lifted over to hg38. We intersected enhancers with enhancers, promoters with promoters, and dyadic elements against both enhancers and promoters to generate a catalogue of high-confidence cCREs.

### Processing 3D Chromatin data

We worked under the assumption that no single method would identify all possible contacts and utilized an ensemble approach, combining methods that work on raw contacts as well as methods that look for areas of contact enrichment.

We downloaded raw sequencing read data for 11 tissues. For the remaining 2 tissues, we used contact data deriving from the tissue type (mammary: HMEC, skin: NHEK). Hi-C contact matrices were generated using the 4DN standardized docker container (v.44)^58^. We employed two methods to normalize and filter adjacency matrices: (1) we used HiCDC+^59^ at false discovery rates of 0.1, 0.01, and 0.001 as inspired from previous work^25^, and (2) we iteratively balanced and applied adaptive coarse-graining to Hi-C matrices before thresholding for top-scoring contacts using cooltools (v.0.7.1)^60^.

We collected or generated loop calls from different methods. To generate DeepLoop calls for K562, HepG2, HMEC, and NHEK, we ran HiCorr and DeepLoop Denoise^61^ at default parameters using normalized in-situ Hi-C alignments from 4DN. For all other models, we downloaded DeepLoop coolers and filtered them by expected / observed ratios to extract the top pixels. We downloaded sample-specific tracks from DeepAnchor^62^: as DeepAnchor calls loops at a resolution of 19bp, we extended each anchor to 5kb total length to match the highest resolution of our Hi-C data. All loops were lifted over to hg38. Peakachu^63^ calls were downloaded from https://3dgenome.fsm.northwestern.edu/downloads/loops-hg38.zip and topologically associating domains were downloaded from https://3dgenome.fsm.northwestern.edu/downloads/hg38.TADs.zip. A full list of chromatin data accessions is available in Supplementary Table S5.

### ChIP-seq peak calling

We collected uniformly processed epigenetic data from EpiMap^57^. For each cell line we downloaded the corresponding epigenomic tracks. For each tissue, we downloaded tracks from 3 different samples, if possible, to generate tracks of averaged signal, prioritizing at least 1 male and 1 female adult track. BigWigs were converted to bedgraphs and lifted over to hg38. If there were multiple samples, the coverage across the bedgraph was average and peaks were called using MACS2^64^ on the averaged bedgraph. A full list of EpiMap accessions is available in Supplementary Table S6.

### CpG methylation

Methylation of CpG sites was downloaded from the ENCODE portal. If multiple bisulfite sequencing experiments existed, we tried to match all tracks to the EpiMap samples and averaged the signal across each bedfile. If not, a single sample was chosen. We considered CpGs as methylated if their percent methylation was >= 80%. For two tissues (NHEK, Hippocampus), methylation data was downloaded from alternative sources but processed with the same methylation percentage cutoff (see Supplementary Table 4).

### Genome-static track processing

To determine RBP binding site clusters, we downloaded binding sites from POSTAR3^65^ and merged any adjacent sites within 10bp. To ensure higher confidence sites, we only keep sites that target multiple target genes and we ensure that the site exists in at least 3 different samples. Additionally, we filter for any sites with the generic annotation “RBP_occupancy”.

Genome-wide maps of recombination rates were downloaded from deCODE^66^. Bigwigs were converted to .wigs and then to bedfiles via BEDOPS (v.2.4.39)^67^. Conserved genomic regions were downloaded from the UCSC phastCons^68^ track and filtered for a conservation cutoff score of 0.7.

We downloaded microsatellite and simple repeat tracks from the UCSC genome browser and reference RepeatMasker^69^ calls from https://repeatmasker.org/genomes/hg38/RepeatMasker-rm405-db20140131/hg38.fa.out.gz. We filtered the RepeatMasker calls to keep “LINE”, “SINE”, “LTR”, and “RNA” repeat classes.

Hot spots were downloaded from Long & Xue^70^. We expanded hot spot clusters based on their individual mutation type and split into separate files for SNPs, indels, and CNVs. Replication hot spots were split into each distinct phase and all data was lifted over from hg19 to hg38.

### Model architecture

The core OGL architecture consists of a rich node feature embedding, graph neural network (GNN) layers for message passing, fully connected layers for representation refinement, and dual task output heads. Because OGLs graph connectivity inherently relies on the underlying input data, we implement a modular architecture that can flexibly adapt to the different construction methods through a joint neural architecture and hyperparameter optimization search space. See Supplementary Note 1 for more information. We describe the most performant architectural details (used for this work) below.

OGL takes as input a graph representation where nodes correspond to genomic elements (genes and regulatory elements) and edges represent chromatin contacts and relationships of genomic proximity. Each node is represented by a feature vector that combines tissue-specific epigenomic features and static genomic properties, which are concatenated with a binned positional encoding to preserve relative genomic positioning information. We partition the genome into 50kb bins according hg38 chromosome start and end points and initialize each bin with a 5-dimensional Xavier uniform embedding^71^. For a genomic feature at chromosome *c* with start position *s* and end position *e*, the positional encoding is computed as:

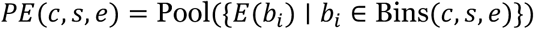

Where 𝐸(𝑏*_i_*) is the embedding vector for bin 𝑏_#_ and Bins(𝑐, 𝑠, 𝑒) returns the set of bin indices that overlap with the genomic interval. If features spanned multiple bins, we used average pooling to combine embeddings. The positional encodings were concatenated to node feature vector as opposed to vector addition to ensure that the initial feature vectors remain interpretable. We also augment Topologically Associating Domains (TADs) to the graphs as “global” nodes acting over a locality to improve expressivity and reflect their role in biology. In this manner, TADs are similar to virtual nodes operating over a smaller local space as opposed to global expressivity.

The most performant OGL configuration uses Graph Attention Network V2 (GATv2) layers which implement dynamic attention. Unlike standard GAT, GATv2 allows the attention function to dynamically adapt to the input features (see^26^). The GATv2 implementation in OGL uses 2 attention heads and the corresponding representations are normalized using GraphNorm^72^. To facilitate gradient flow, we implement distinct-source residual connections. For node representations ℎ*_l_* 𝑎𝑛𝑑 each layer 𝑙, the residual connection is computed as

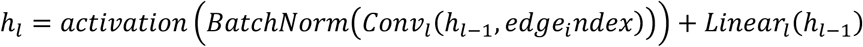

where Linear is a learnable projection. The node representations from graph convolutional layers are further processed through fully connected layers to refine feature extraction. Each fully connected layer is followed by a LayerNorm operation and GELU activation:

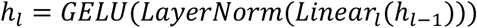

Additionally, dropout with rate is applied after each fully-connected layer.

OGL employs a multi-task approach to simultaneously predict regression and classification. The regression task head predicts log2 TPM values for gene expression while the classification task head performs binary classification to determine whether gene expression is high or low. The task heads compute:

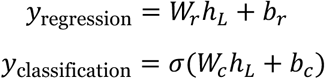

where ℎ*_L_* is the final node representation, 𝑊*_r_* and 𝑊*_c_* are learnable weight matrices, 𝑏*_r_* and 𝑏*_c_* are bias terms, and σ is the sigmoid activation function. The model loss is a weighted combination of a regression loss and binary cross-entropy loss for classification, where expression classes are high (> 0 log2 TPM) or low (<= 0 Log2 TPM) based on the bimodal distribution of expression targets. The combination loss function is defined as:

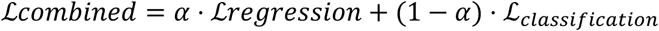

where α is a hyperparameter that controls the relative importance of the regression task.

### Model training

Our model specifically focuses on the expression of protein coding genes which we split via whole-chromosome holdouts to prevent data leakage and regulatory overlap. We reserved all genes on chromosome 8 and 9 for testing, and all genes on chromosome 10 for validation. All other protein coding genes were used for training. Cell line models regressed gene-level log_2_ TPM values derived from bulk RNA-seq. Given the redundancy of tissue samples across the EpiMap repository, we chose to combine tissue-average tracks during pre-processing and aimed to build “tissue-general” models that regress the median gene-level log_2_ TPM values of that gene within that tissue in the GTEx V8 dataset. We employ dual masking during training: a propagation mask allows the network to update representations for all nodes in the forward pass, while a separate loss mask ensures that only designated gene nodes contribute to the loss computation and subsequent gradient updates during backpropagation. This approach enables the model to leverage the full graph structure for representation learning while focusing optimization specifically on gene prediction targets.

All node features were scaled twice: first with a quantile scaler to remove the effects of outliers before standardizing values between 0 and 1. Scaling was applied first to the training set before the same factors were applied to the test and validation sets. All models were constructed in PyTorch Geometric (v.2.5.3)^73^. Optuna (v.4.0.0)^74^ was used for hyperparameter optimization on a subset of chromosomes against validation RMSE. Best models utilized Smooth L1 as the regression loss, an α weighing factor of 0.95 for the combination loss function, batch size of 16, learning rate of 5e^−4^, the AdamW^75^ optimizer, distinct source skip connections, GATv2 operator with 2 attention heads, 2 convolutional graph layers and 2 fully connected layers with a model width of 200. Models were trained for 60 epochs with an early stopping patience of 12 where the registered best validation was smoothed over the last 5 epochs. We used a cosine learning rate scheduler and employed warm-up where we linearly increased the learning rate from 0 to the target value over 10% of the training steps. For regularization, weight decay of 0.02 is applied alongside dropout *p*=0.1. All models were run three times across three random seeds.

### Saliency and attention weights

Given a GNN model with loss function 𝐿, the raw gradient saliency attribution scores gradient saliency attribution scores 𝑠 for node features 𝑥 are computed as:

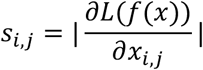

Where 𝑖 indexes the nodes in the graph and 𝑗 indexes the feature dimensions. For visualization, we normalize the saliency values using percentile-based scaling. For each feature dimension, we compute the 5^th^ (𝑞*_low_*) and 95^th^ (𝑞*_high_*) percentile of the saliency values. We then normalize each value using the formula:

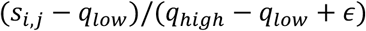

Finally, we apply a clamp operation to restrict all values between 0 and 1.

For each edge (𝑖, 𝑗) in the graph, we compute the average attention weight 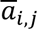 by the formula:

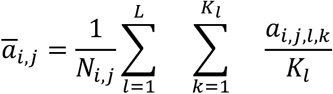

Where 𝑎*_i,j,l,k_* is the attention weight for edge (𝑖, 𝑗) in layer 𝑙 and head 𝑘, 𝐾*_l_* is the number of attention heads in layer 𝑙, 𝐿 is the total number of layers, and 𝑁_𝑖,𝑗_ is the number of times the edge (𝑖, 𝑗) appears across all batches. This approach first averages attention values across heads within each layer, then aggregates these means across all occurrences of the edge in different batches to produce a single representative attention weight for each edge in the graph.

### *In-silico* experiments

Perturbation and interpretability experiments were run on the entirety of each protein-coding gene’s *K* = 2 graph structure with 2 layers chosen commensurate to the number of convolutional layers in the model.

For each perturbation, we calculate the relative fold change between each baseline prediction and the effect as:

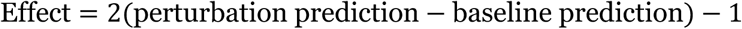

### Subgraph-level feature ablations

We quantified the impact of each feature by measuring differences between baseline and perturbed model predictions. For each protein-coding gene node, we extracted its *K*-hop subgraph, and for each feature *f* and node *n*, we computed:

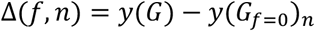

Where 𝑦(𝐺) represents the model’s prediction on the original subgraph and 𝑦(𝐺*_f_*_=0_)*_n_* is the prediction at node 𝑛 when feature 𝑓 is set to zero across the entire subgraph in the pre-scaled space.

### Node deletions

To focus on accurate predictions, we first binned genes into high (≥ 5 TPM), medium (5 > TPM ≥ 1), and low (1 > TPM > 0) log_2_ expression values. For high and medium bins, we kept all genes with ≤ 0.05 TPM difference between predicted and observed. For the low bin, we applied the same criteria but capped selection at 1000. We then quantified the impact of each individual node by measuring the expression change caused by their removal. For each gene *g* and neighboring node *n*, we computed:

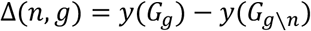

Where 𝑦(𝐺*_g_*) represents the model’s prediction on gene 𝑔*’*s original *K*-hop subgraph, and 𝑦G𝐺*_g_*_∖*n*_ K is the prediction after removing node 𝑛.

### Node-level feature ablations

We then categorized nodes by their regulatory class (enhancer, promoter, dyadic element, or gene) and ranked them within each class by their predicted impact from the node deletion experiment. We extracted the top 2,500 most impactful nodes from each regulatory class, resulting in 10,000 high-impact nodes per tissue or cell line. For each of these nodes, we extracted the *K*-hop subgraph (*K*=2) containing the node and its connected gene, then systematically perturbed each feature by setting its value equivalent to 0 in the original feature scaling space.

Formally, we implemented a targeted ablation approach where for each node-gene pair, we computed the feature-specific relative fold change:

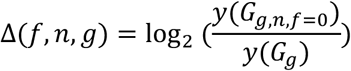

Where 𝑦(𝐺*_g_*) represents the model’s prediction on gene 𝑔’s original *K*-hop subgraph, and 𝑦(𝐺_(*g,n,f*=0)_) is the prediction when feature 𝑓 of node 𝑛 is set to zero.

### Feature interaction analysis

We computed partial correlation matrices for all possible feature pairs across perturbed elements to generate a correlation matrix where interactions are conditioned against every other feature. For each element, we identified features with significant perturbation effects (absolute relative fold change ≥ 0.1). Pairs of these features were classified as *synergistic* if both features impacted expression in the same direction (both positive or both negative) and *antagonistic* if they impacted expression in opposite directions. We then aggregated these interaction counts across all samples to identify recurring partners.

## Extended Data

**Extended Data Fig. 1.**
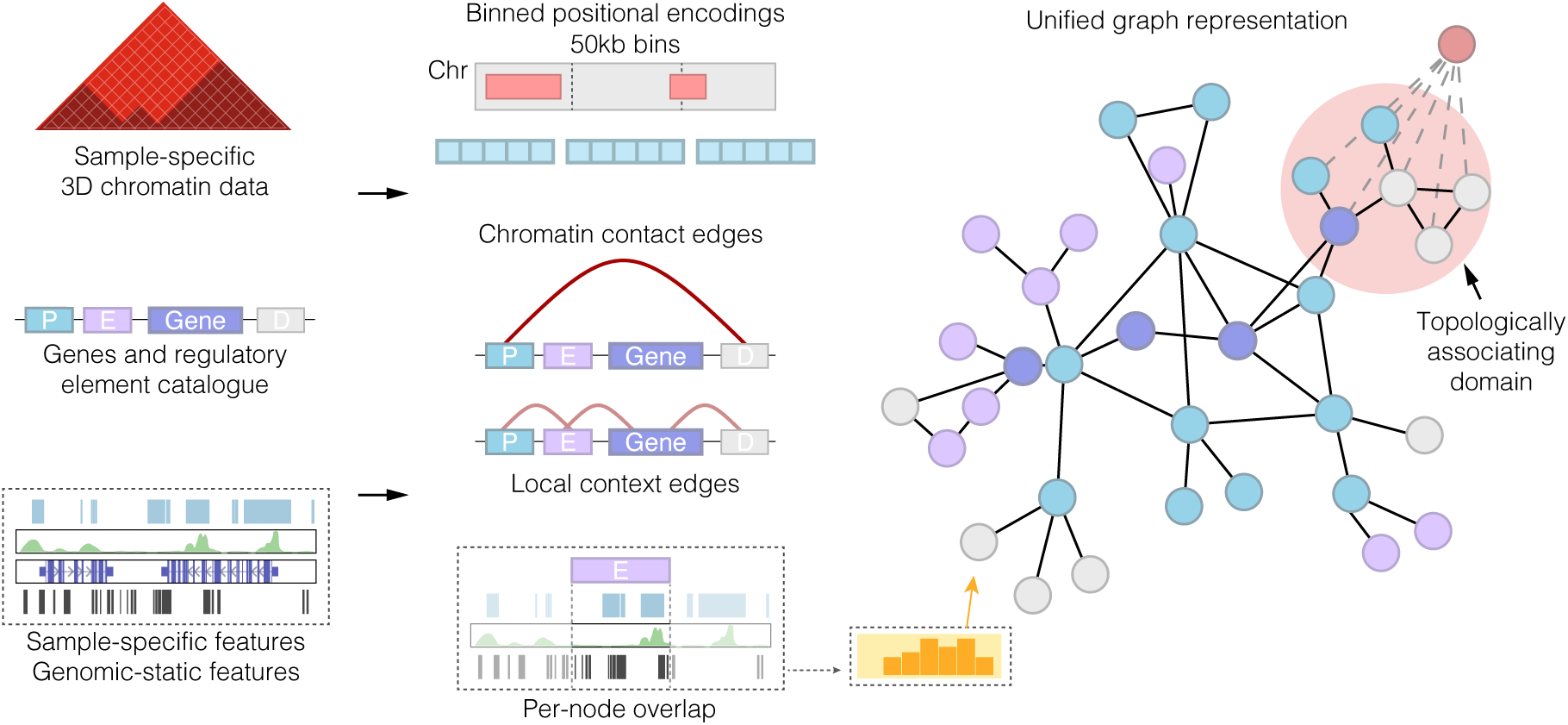
Overview of OGL’s graph construction. OGL takes 3D chromatin data, gene annotations, a regulatory element catalogue, and a series of data tracks as input. Edges are drawn between nodes based on overlap with chromatin anchors and presence within a local context window. Binned position encodings get added to the feature vector to inform the model of relative positioning. Topologically associating domains connected all nodes within its boundaries, acting as a “local global” node to increase expressivity.

**Extended Data Fig. 2.**
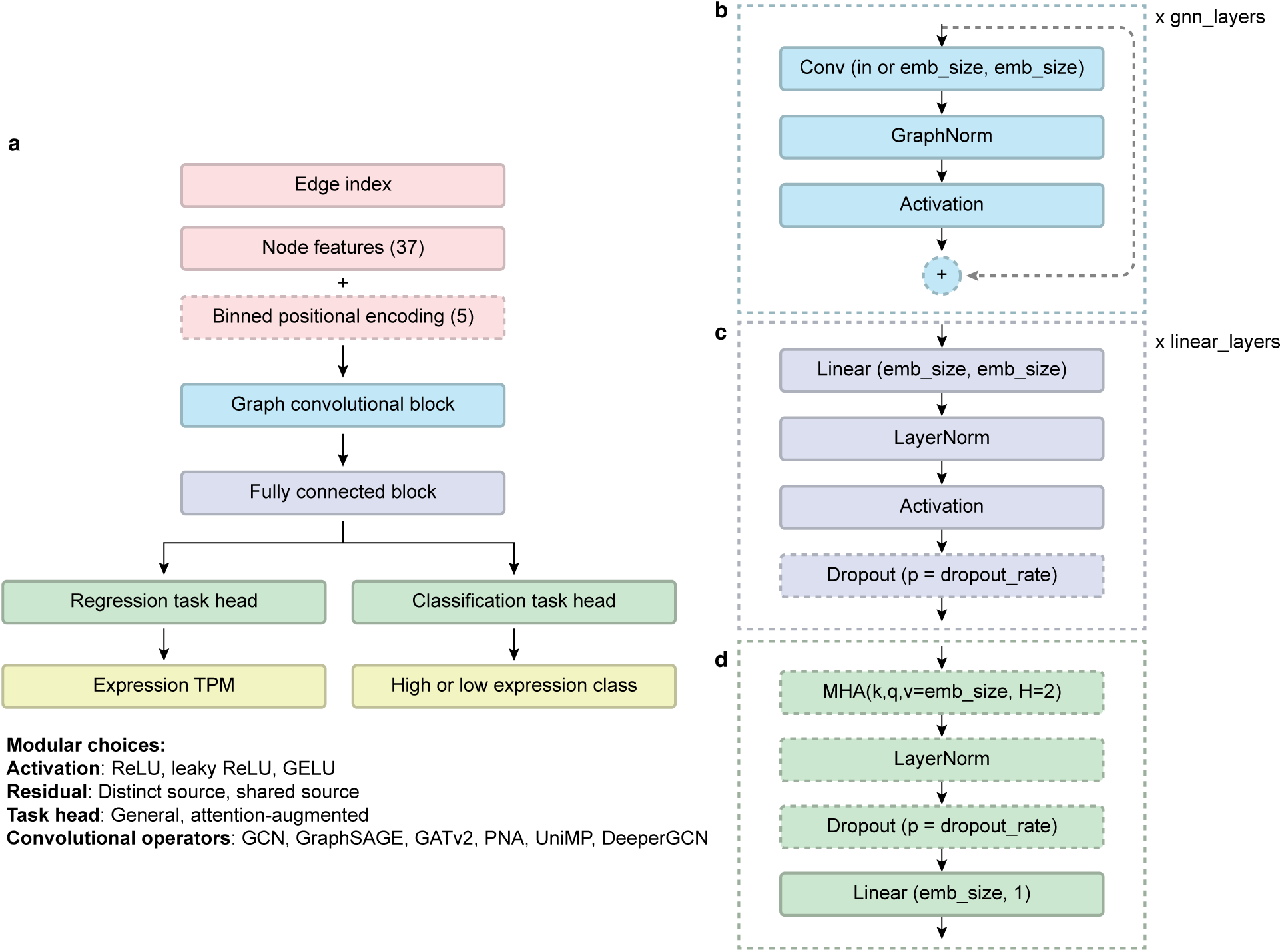
Modular GNN architecture of OGL. a, OGL-pipeline features a highly modular and flexible architecture solution space that can flexibly adapt to the graphs via a joint optimization process. a, The general multi-task GNN architecture with modular components listed. b, Definition of different network blocks that comprise the neural network layers. Dashed boxes indicate optional components.

**Extended Data Fig. 3.**
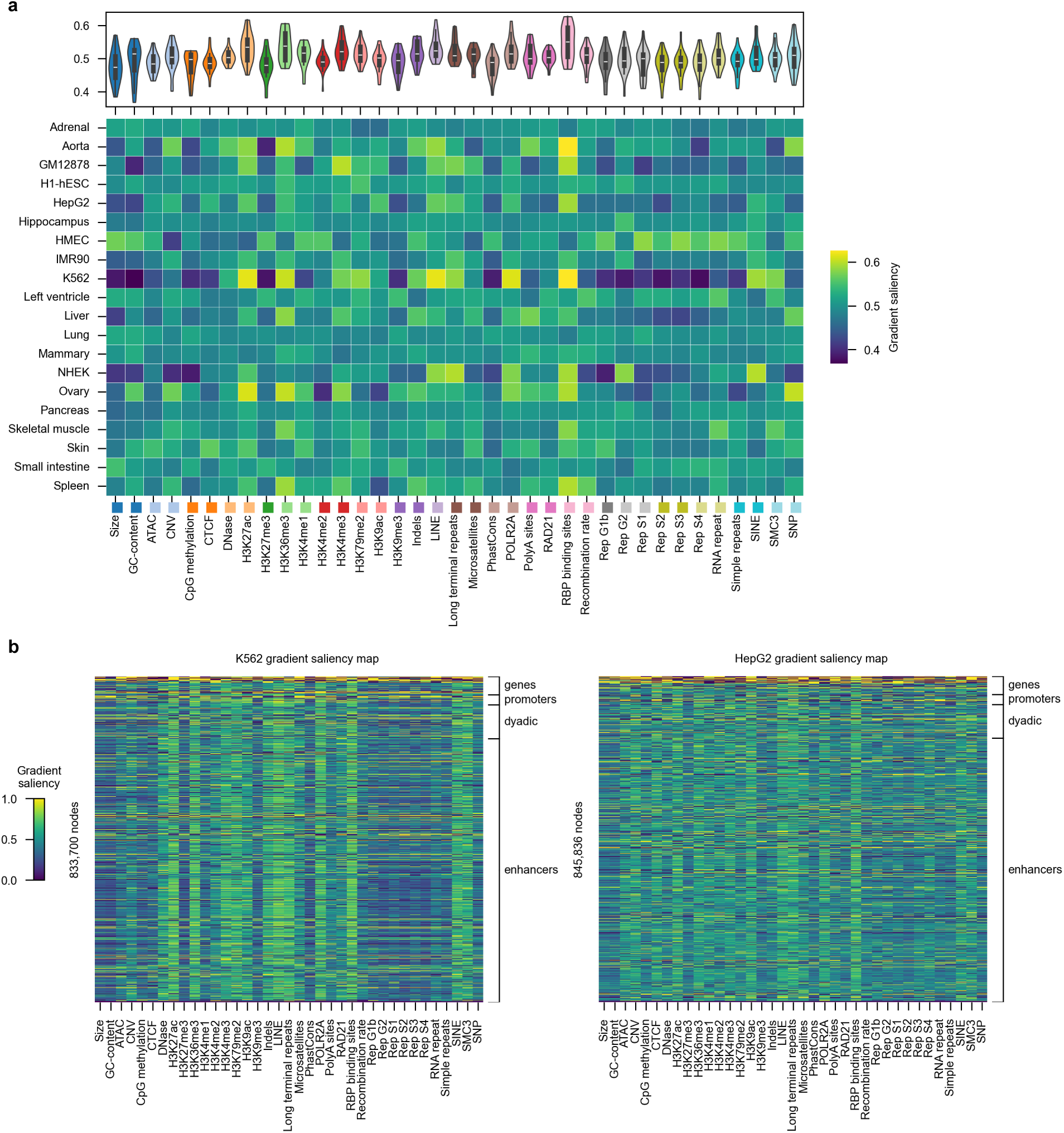
Gradient saliency maps show sample-specific sensitivity of features. a, Heatmap of the averaged saliency gradients per node feature. b, Saliency map across nodes of K562 model shows high sensitivity across specific features such as H3K27ac and RBP binding sites whereas saliency map across nodes of HepG2 show lower gradients at individual features and more consistent sensitivity across the feature space.

**Extended Data Fig. 4.**
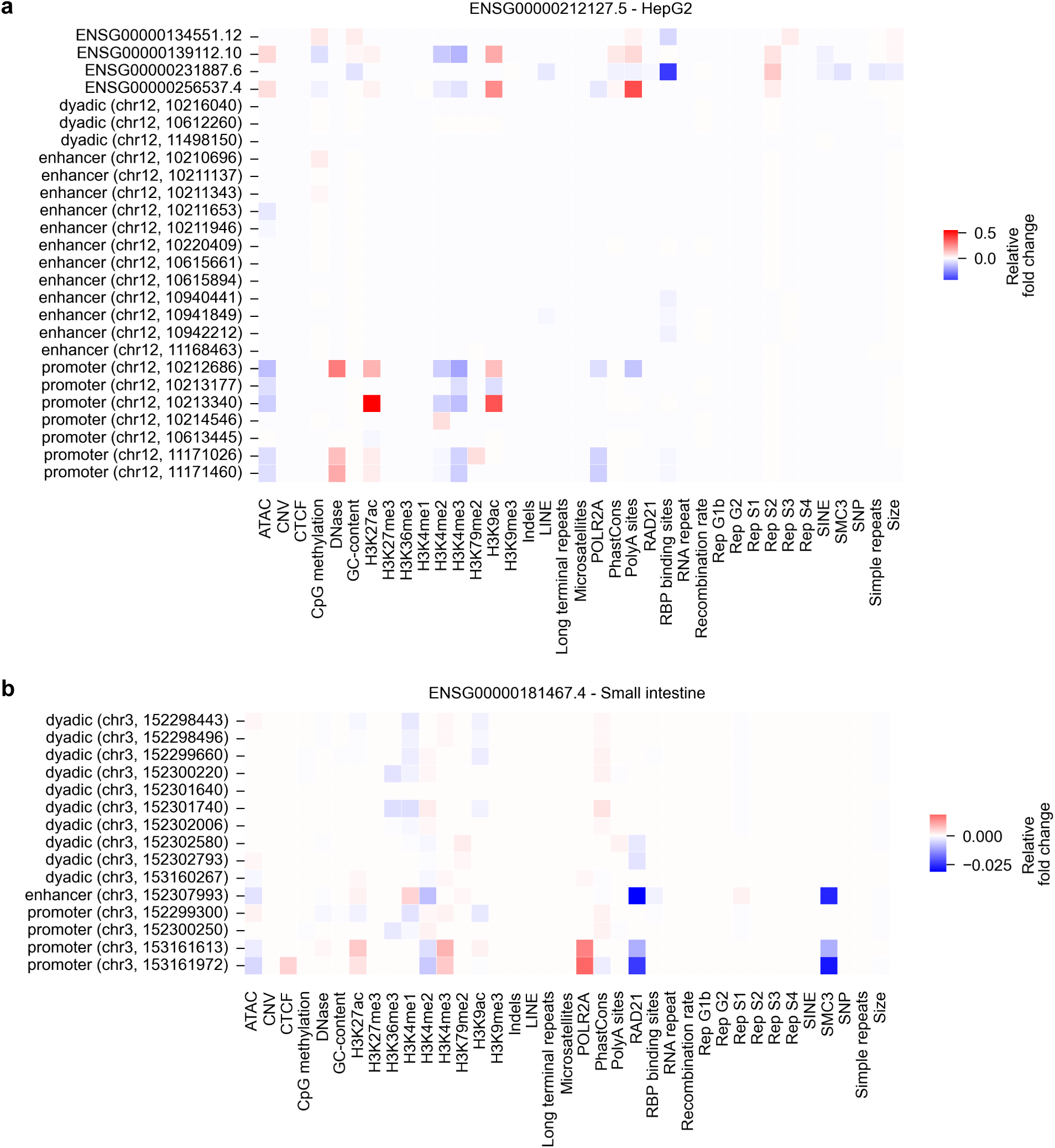
Node-level feature perturbations reveal varied feature utilization by elements connected to genes. Feature ablation effect at the scale of individual features within an element, demonstrating the granularity of OGL’s multi-scale perturbation framework. a, Feature ablation effects for elements regulating ENSG00000212127 (TAS2R14, HepG2). b, Feature ablation effects for elements regulating ENSG00000181467 (RAP2B, small intestine).

**Extended Data Fig. 5.**
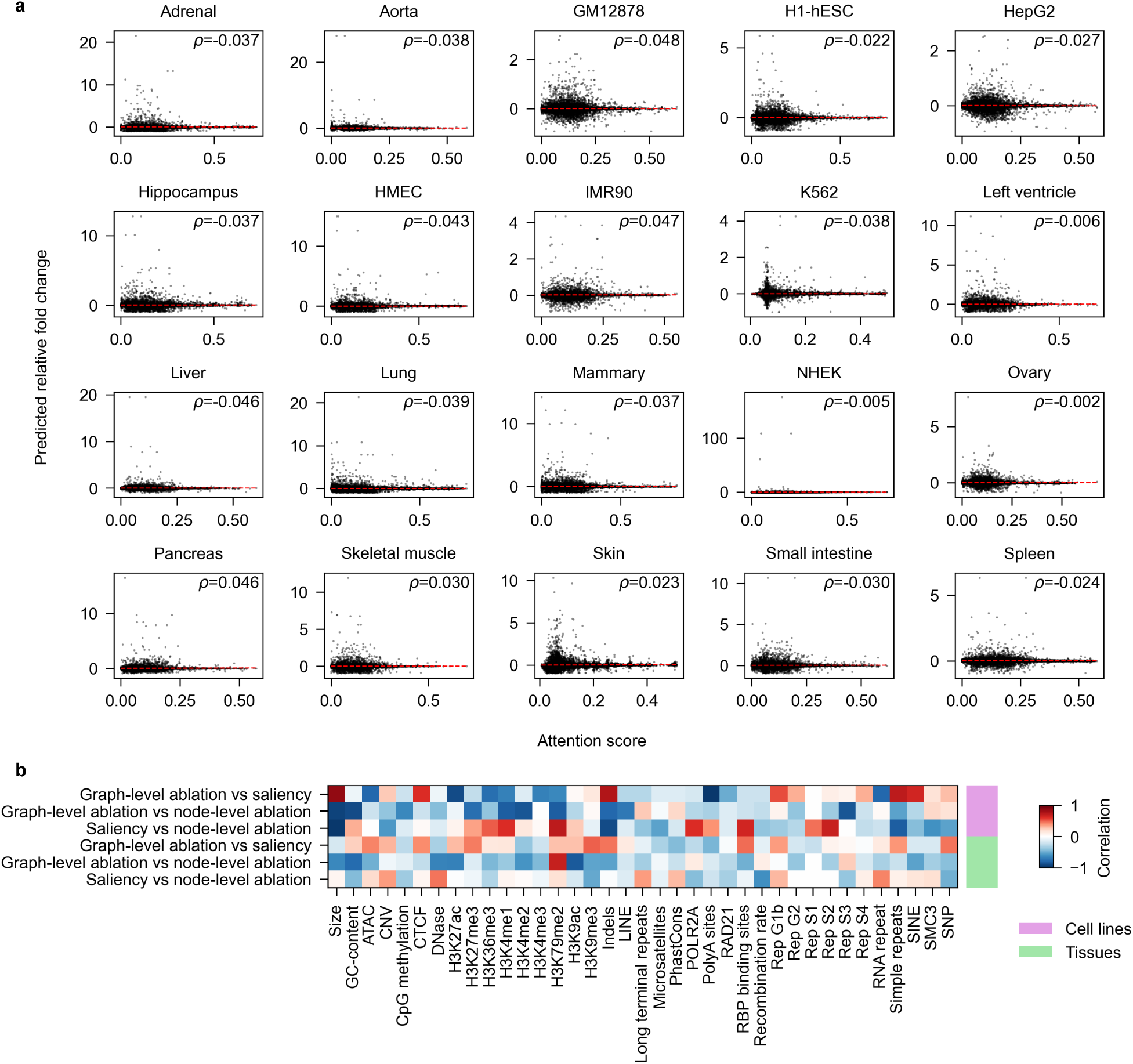
Correlation between interpretability methods reveals challenges in interpreting deep neural networks. a, Scatter plots comparing perturbation predictions against attention score, showing little correlation across all models. b, Correlation between different interpretability methods across the feature space. “Graph-level ablation” refers to full feature ablation. “Saliency” refers to integrated gradient contribution score. “Node-level ablation” refers to ablation of individual features at the node level.

## Notes

https://github.com/sciencesteveho/ogl

### Competing Interest Statement

The authors have declared no competing interest.

