## Supplementary Information for "Contextualizing gene expression with feature rich graph neural networks"

**Supplementary note 1:** Design philosophy of OGL and its pipeline
**Supplementary note 2:** GNN “solution space”
**Supplementary note 3:** Hyperparameter optimization and neural architecture search
**Supplementary note 4:** Graph construction methods
**Supplementary note 5:** Saliency methods
**Supplementary note 6:** Gene-set enrichment analysis
**Supplementary note 7:** CRISPRi validation
**Supplementary note 8:** Additional multi-omics pre-processing
**Supplementary note 9:** Tissue-specificity analysis

**Supplementary Fig. 1:** Relationship between model performance and number of chromatin contacts
**Supplementary Fig. 2:** K562 element-specific saliency maps
**Supplementary Fig. 3:** Comparison of saliency for high sensitivity nodes in both HMEC and IMR90 models
**Supplementary Fig. 4:** Gene type annotation of high sensitivity genes
**Supplementary Fig. 5:** Sensitivity of experimentally validated cCREs
**Supplementary Fig. 6:** Predictive performance of models trained with randomized features
**Supplementary Fig. 7:** Joint feature ablations reveal non-additivity
**Supplementary Fig. 8:** Gene set enrichment analyses of most affected genes from full H3K27me3 ablation
**Supplementary Fig. 9:** Sample-specific gene-set enrichment of most affected genes from full node feature ablation
**Supplementary Fig. 10:** Tissue-specificity of node perturbation effects.
**Supplementary Fig. 11:** Partial correlations of feature effects reveal the cell-type differences in regulation.
**Supplementary Fig. 12:** Optimization showing convergence over 200 trials
**Supplementary Fig. 13:** OGL’s end-to-end design facilitates rapid iteration

### Supplementary note 1: Design philosophy for OGL and its pipeline

The application of deep learning methodologies and diverse neural architectures – including convolutional neural networks (CNNs)[1–4], transformers[5,6], and graph neural networks (GNNs)[7,8] – has revolutionized genomic analysis, enabling significant advancements in regulatory motif identification[9], variant effect prediction[10], gene expression modeling[6], and more[11,12]. While frameworks such as PyTorch[13], TensorFlow[14], and JAX[15] have substantially reduced implementation complexity through abstraction, the development of effective deep learning genomic models remains a multifaceted challenge requiring integrated expertise across genomics, computer science, and machine learning domains. This process necessitates systematic execution of several critical phases, including: (1) data preparation and preprocessing, (2) model architecture and design, (3) training and hyperparameter optimization, (4) evaluation and deployment.

Packages such as Selene[16], gReLu[17], EUGENe[18], ENNGene[19], and AMBER[20] aim to decrease the implementation barrier and significantly decrease the effort required to implement genomic deep learning models. Other tools, such as TF-MoDISco[21], fastISM[22], and CRÈME[23] decrease the implementation barrier for model interpretation. However, these platforms dominantly focus on CNN implementations, leaving alternative architectural paradigms underexplored.

GNNs represent a class of neural architectures that explicitly model relational inductive bias through node-to-edge connectivity patterns. Unlike CNNs which leverage a spatial inductive bias, or transformers which leverage a global inductive bias between inputs, GNNs directly encode relationships between entities through graph structure. Their applications to genomics are relatively new and primarily employ a binning approach where genomic loci are discretized into fixed-size nodes which are connected via 3D chromatin interaction data[7,8,24]. This approach presents two fundamental limitations: (1) binning necessitates the fragmentation or aggregation of genomic features, resulting in distorted representations of biological entities, and (2) the resulting models are ill-suited for *in-silico* perturbation studies due to the entanglement of multiple regulatory elements within individual bins.

We propose an alternative modeling paradigm wherein individual nodes correspond to complete genomic elements, analogous to gene regulatory networks where each node represents an entire gene. This approach generates representations specific to biologically meaningful elements and inherently facilitates systematic *in-silico* perturbation analysis. However, the literature provides limited precedent for developing such models, raising numerous methodological questions spanning graph construction, feature selection, and architectural design. Critical considerations include which molecular features and regulatory element classes to incorporate, which graph convolutional operators to employ, optimal network depth, and whether to model tissue-specific or cross-tissue relationships.

Identifying optimal architectural configurations for novel modeling paradigms presents a significant challenge. This challenge is compounded by the universal approximation theorem[25–28], which establishes that neural networks of sufficient complexity can theoretically approximate any continuous function. Empirical evidence further demonstrates that architectural superiority is highly context-dependent and often counterintuitive as highlighted by several studies:

1. In Tönshoff et al.,[29] simple fine-tuned graph convolutional networks (GCNs) with increased width and residual connections outperformed transformer based models.
2. In the Open Graph Benchmark Large Scale Challenge (OGBLSC), Veličković et al.[30] demonstrated that deep GCNs achieve start-of-the-art performance with minimal architectural modifications.
3. Huang et al.[31], showed that label propagation combined with multi-layer perceptrons surpassed sophisticated GNN implementations.

4. In the OGB-products leaderboard, vanilla GraphSAGE achieves 96% of the test-set accuracy of the leading model.
5. In Luo et al.[32], minimally alerted architectures achieve 95% of the test-set accuracy of loading models compared to the OGB-proteins leaderboard and classic GNN operators are as performant as graph Transformers.

Similar results exist outside of graph-based work[33–35], including a recent DREAM competition that saw a variety of architectures in the top performing models[36]. These findings align with the "no free lunch theorem": no universal architectural solution exists across all problem domains[37].

Empirical approaches to architecture and hyperparameter selection are essential considering the difficulty of *a priori* architectural optimization and comprehensive empirical evaluation is critical when novel modeling paradigms lack historical precedent. Without established heuristics to guide architectural decisions, systematic exploration of the design space through rigorous optimization frameworks represents the most viable path toward identifying performant model configurations.

To address these challenges, we developed a modular, end-to-end pipeline designed to: (1) automate the construction of omics-based graph representations, (2) facilitate rapid experimental iteration, and (3) perform joint neural architecture and hyperparameter optimization to efficiently identify performant configurations. In terms of our final architecture (best performing OGL models), we utilized a two-step hybrid approach: first, we perform neural-architecture search on a subset to lock in key parameters, then we fine-tune the parameters on the full dataset and make targeted modifications to arrive at high performing architectures.

### Supplementary note 2: GNN "solution space"

We adopt a GNN "solution space" analogous to the design space of GraphGym as implemented by You, Ying, and Leskovec[38]. Given the breadth of the graph construction, we aimed for a modular solution space so that the GNN architecture could flexibly adapt to a diverse range of graph construction choices. Each graph inherits from a "ModularGNN" class of the general form: input, operatory layers, fully-connected layers, task head, output. Models span three classes of GNN architectures:

(1) Classic message passaging neural networks (GCN[39], GraphSAGE[40], PNA[41])
(2) Attention-based models (GATv2[42], UniMPTransformer[43])
(3) Large-scale models (DeeperGCN[44])

Other modular aspects of the models, including network depth, width, activation function, and residual connections are listed in Extended Data Fig. 2.

### Supplementary note 3: Hyperparameter optimization and neural architecture search

We run a distributed instance of a joint hyperparameter and architecture search via Optuna[45]. Since neural architecture search is generally very computationally expensive, we perform our search on a subset of 12 chromosomes (60%) as a way to converge onto a baseline strong-performing architecture that can be tuned while avoiding overfitting to the dataset. Trials were pruned if their validation performance fell below the median of the intermediate results. Results of our architecture search are presented in Supplementary Fig. 12.

### Supplementary note 4: Graph construction methods

OGL creates omics-based graphs based on user-specified node types (Supplementary Table 1) and creates edges based on two sources of information: (1) the locality of a genomic locus, or "local context", and (2) the distal interactions of a genomic locus as informed by 3D chromatin edges. Users specify and input "feature window" and for any given node, edges are created between that node and all other nodes in the feature window. Users also specify a 3D chromatin contact file, which can be any chromatin contact assay in .bedpe format. Edges are created between regulatory node types if the nodes fall within a window to a chromatin contact anchor, which we specify as a conservative 1,500bp. Edges are created to gene nodes if chromatin contacts fall within 2kb of an annotated transcription start site from RefTSS v4[46]. Additional interaction data types, specifically micro-RNA interaction networks and RNA binding protein interaction networks can be optionally appended to the 3D interaction graph. Node and edge connections from both the local context window and the 3D interactions are combined to create the full graph view.

For the implementation of this paper, we specifically used a local context of 12.5kb, corresponding to the median compartment length discovered in Harris et al.[47]

Graphs undergo further processing to be fit for training. We prune all nodes that overlap the ENCODE v2 blacklist[48], isolated nodes, self-loops, and nodes that never hop to genes as they will not inform the latent representations. We append each node with an optional 5-dimensional binned positional encoding and a 37-dimensional feature vector derived from each node's overlap against a mix of tissue-specific and genome-static datasets including replication hot-spots, reference repeat annotations, and histone modifications.

It is pertinent to note that graph construction is heavily influenced by the underlying input data. For example, the resolution of chromatin contacts directly impacts the centrality and connectivity of each graph, as does the choice of regulatory catalogue (which varies with size and location), the size of the local context window (which directly increases the number of edges created), and the selection of different node types and interactions to append to the base graph. Thus, the created graphs are highly sensitive to changes in graph construction parameters, affirming our reasoning for employing Neural Architecture Search (NAS). NAS allows us to converge on performant architectures that flexibly adapt to a wide variety of different graph construction methods.

We ran a series of experiments varying graph construction, such as using different regulatory element catalogues, interaction types, and adding additional node representations. We ultimately settled on the choices of the main paper for a balance of computationally efficiency, interpretability, and performance stability. A subset of graph construction experiments are documented in Supplementary Fig. 13.

### Supplementary note 5: Saliency methods

In principle, gradient × input methods are often favored for measuring first-order contributions of features to model predictions. However, in our setting, where numerous node features are sparse or near zero and are distributed across multiple molecular annotations, raw gradient values often yield clearer, more interpretable saliency signals. Specifically, input × gradient tends to diminish saliency for features that have very small magnitudes, even if a small increase in those features could substantially shift the model's output. For example, with input × gradient, zero-valued inputs will always have zero attribution (0 x anything = 0), thereby providing little information for sparse inputs. By contrast, raw gradients directly capture the potential impact of small perturbations in each feature, making them especially helpful for identifying which features or nodes the model is most sensitive to. In our analysis of OGL models, we rely on raw gradients as a pragmatic measure of local sensitivity in contexts where features are often smaller and can vary widely in their scaled values compared to the one-hot encodings of sequence-to-function models

We emphasize that these raw gradient scores do not represent absolute “feature contributions” but rather a measure of how much small changes in each feature would alter the model’s prediction. We found this perspective aligns well with our broader aim of identifying features that have high potential to impact regulatory activity. Thus, while gradient × input remains a valuable tool, raw gradients here provide a useful metric of each feature’s capacity to influence predictions, making it well-suited for our feature-rich GNN architecture.

We note that for the comparisons between saliency and perturbations in Extended Data Fig. 5, we employ integrated gradients as our comparative framework due to their fundamental axiomatic properties. This method integrates gradients along a path from a baseline to the input, addressing the zero-attribution issue while maintaining attribution completeness. Thus, integrated gradients are an appropriate and effective benchmark for evaluating the performance of different attribution approaches over raw saliency.

### Supplementary note 6: Gene-set enrichment analysis

We used Enrichr[49] to run gene-set enrichment analysis through GSEApy[50] (v.1.1.5) with the “GO Biological Process 2023”, “GO Molecular Function 2023,” and “Reactome Pathways 2024” gene sets.

### Supplementary note 7: CRISPRi validation

While previous work using CRISPRi to validate neural network models have focus on individual datasets, we specifically utilize curated gold-standard CRE-gene interactions from Gschwind et al.[51]. The authors collect CRISPRi CRE-gene pairs across multiple studies and thoroughly investigate the statistical power of the CRISPRi experiments.

Our initial investigation of CRISPRi effects revealed a mean effect size of approximately 1%. For context, we calculated the expected fold change between two replicates of RNA-seq in K562 to be 4.33% (accessions ENCFF384BFE and ENCFF611MXW). Given these modest effect sizes for most cCRE-gene links, focusing on the entire dataset would likely yield inconclusive results. Therefore, we established a filtering threshold of ≥10% for CRISPRi effects to focus specifically on high-impact CRE-gene interactions that could be reliably distinguished from experimental noise. We retained enhancer-gene links where the gold-standard enhancer loci overlapped with our regulatory catalogue and connected to the same gene in its $K$-hop subgraph.

### Supplementary note 8: Additional multi-omics pre-processing

#### Micro-RNA processing

All Homo sapiens targets for miRNAs were downloaded from MirTarBase[52,53] and filtered for functional evidence. We then derived miRNA coordinates and miRNA targets from the filtered catalogue.

Active miRNAs were determined from microRNA-seq datasets. Gene quantifications (miRNA) were downloaded from ENCODE and we matched biosamples when possible. We normalized counts to CPM (counts per million) and considered any miRNA with > 5 CPM as active. We removed miRNAs from our genome annotations as to avoid training our model to learn their expression – given the direction of the central dogma, though transcriptional efficiency is not 100%, there is still the implication that an active miRNA implies expression of the miRNA gene – and thus removing miRNA genes allows us to avoid circular data where the presence of a feature implies its own outcome.

#### RNA-binding protein network

To create the RNA binding protein network, we intersect the location of RBP binding sites downloaded from POSTAR3[54] with the GENCODE v26 annotation. Any direct overlaps of a binding motif with gene annotation were considered a link from RBP to gene. We only keep RBP binding sites with a specific RBP and not the generic annotation "rbp_occupancy". Additionally, we only kept RBP binding sites present in at least 3 separate samples. If the RBP network was added to the graph for training, the RBP genes in the network were removed from the regression task.

#### Calling cis-regulatory modules

To call CRMs, we use peakMerge.py from ReMap2022[55] on our epigenomic tracks described above. We use all narrow peaks with cutoff of 5 as input to enforce a stricter FDR for the cis-regulatory modules.

### Supplementary note 9: Tissue specificity analysis

We calculate Tau scores for nodes based on their max and average exerted effects across samples. For each element, we first normalized its perturbation effects across tissues by dividing by the maximum absolute effect and calculated Tau as the sum of:

$$\frac{(1 - \text{normalized effects})}{(\text{number of tissues} - 1)}$$

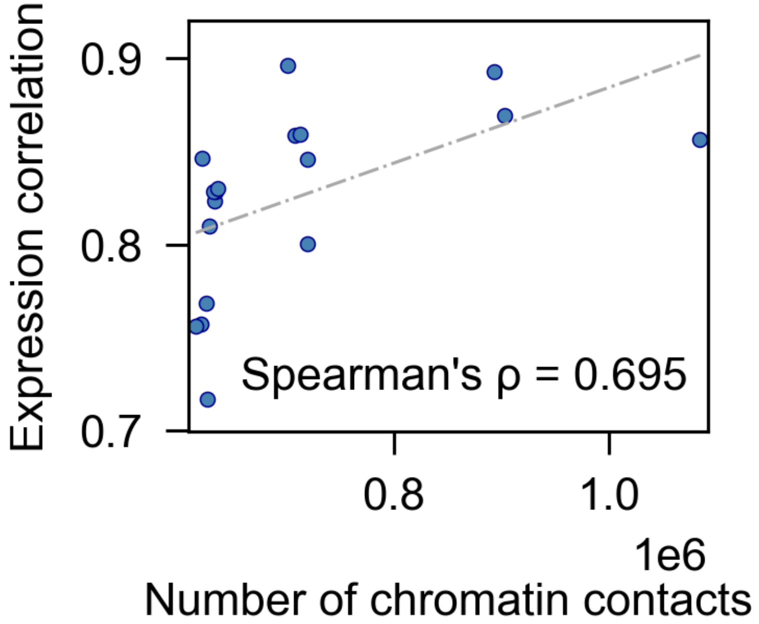


**Supplementary Fig. 1 | Relationship between model performance and number of chromatin contacts.** Each scatter point represents an individual model. X-axis shows total amounts of chromatin contacts used for the model, and Y-axis shows Pearson correlation on hold-out test genes.

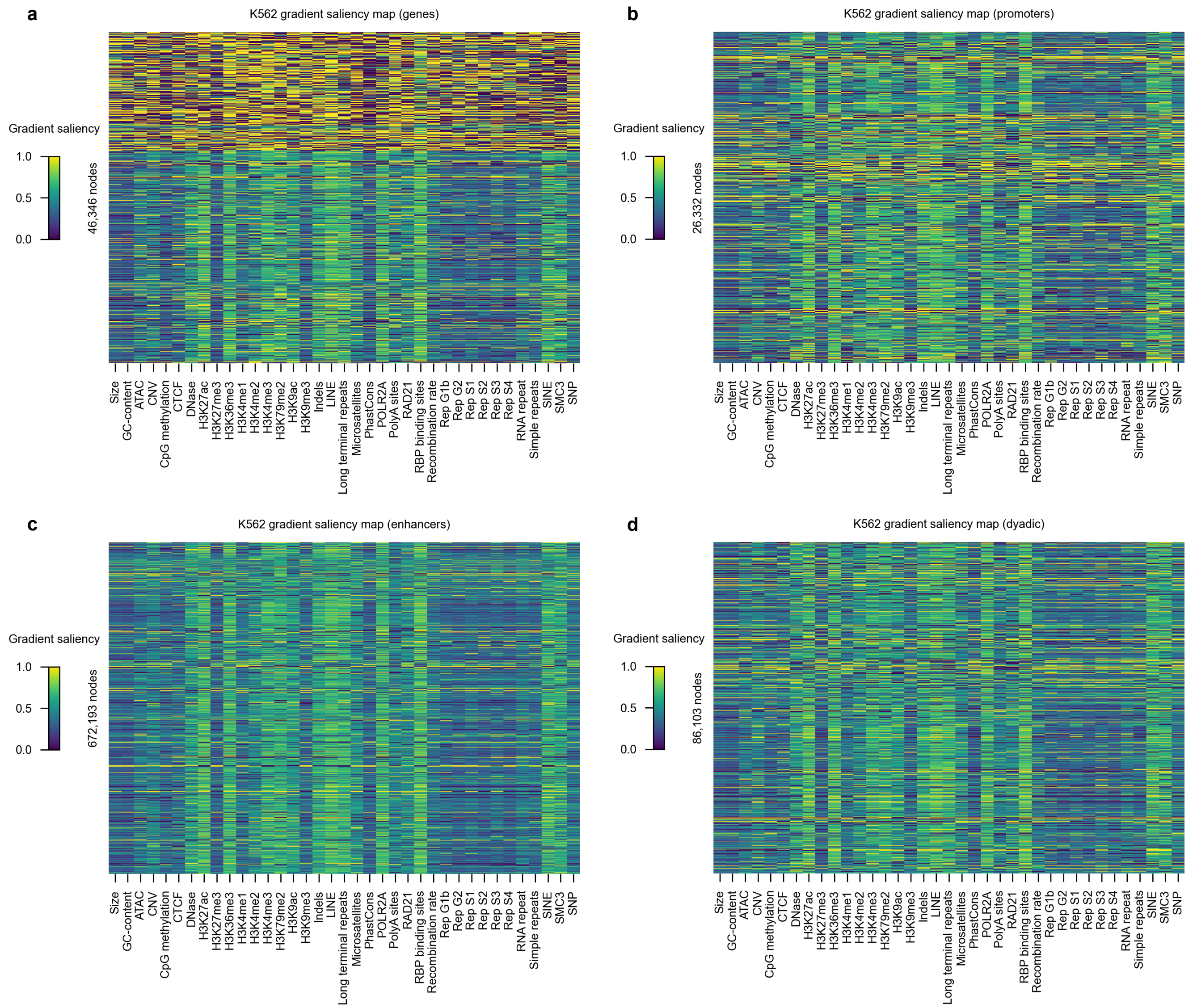


**Supplementary Fig. 2 | K562 element-specific saliency maps.** Zoomed-in saliency maps showing feature sensitivity for **a**, genes, **b**, promoters, **c**, enhancers, and **d**, dyadic elements.

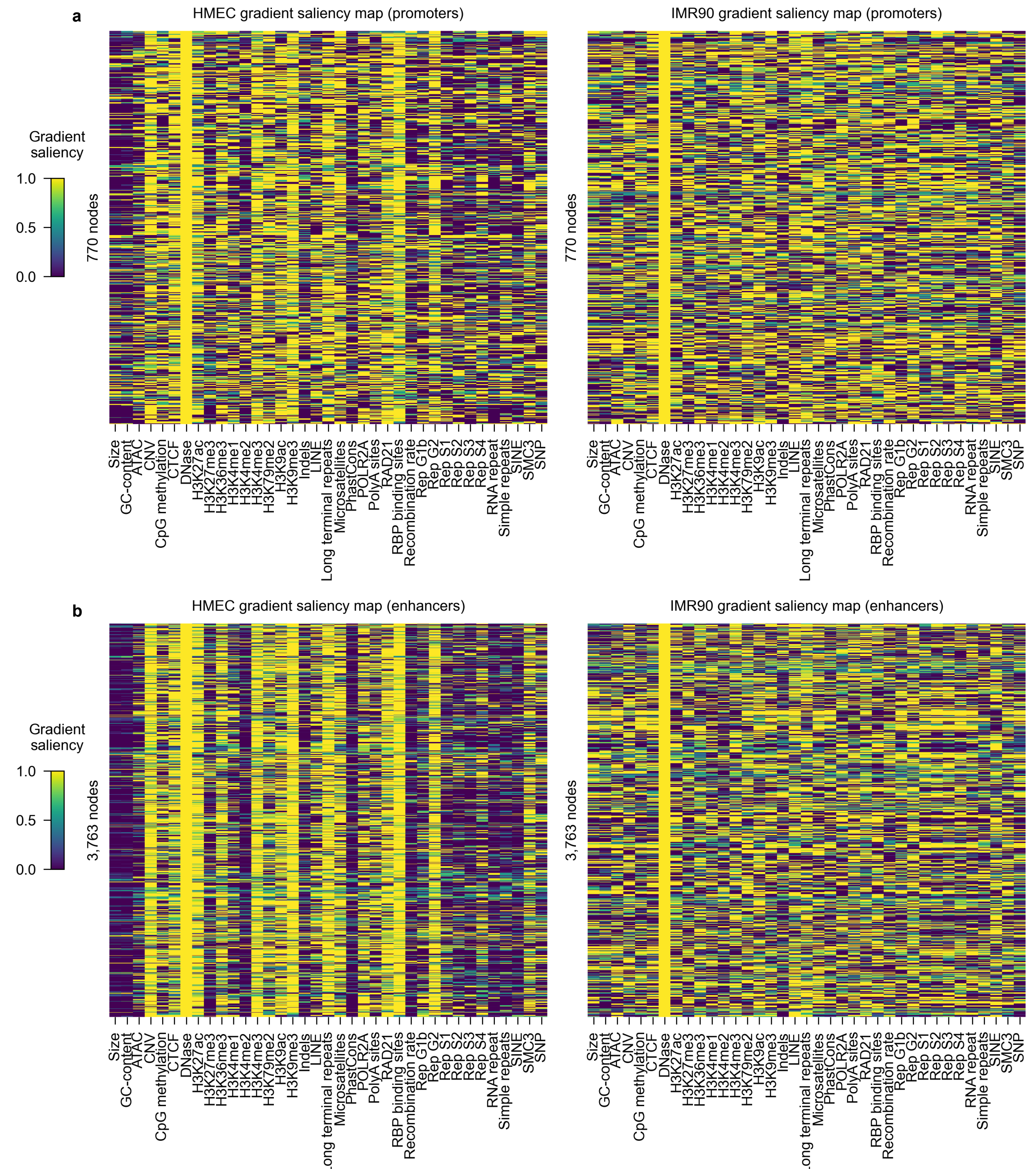


**Supplementary Fig. 3 | Comparison of saliency for high sensitivity nodes in both HMEC and IMR90 models.** High-sensitivity enhancers and promoters present in both HMEC and IMR90 models filtered for scores of 1 (max gradient) according to the DNase track. The saliency map of the enhancers (**a**) shows a larger difference in sample-specific patterns compared to the differences between promoters (**b**).

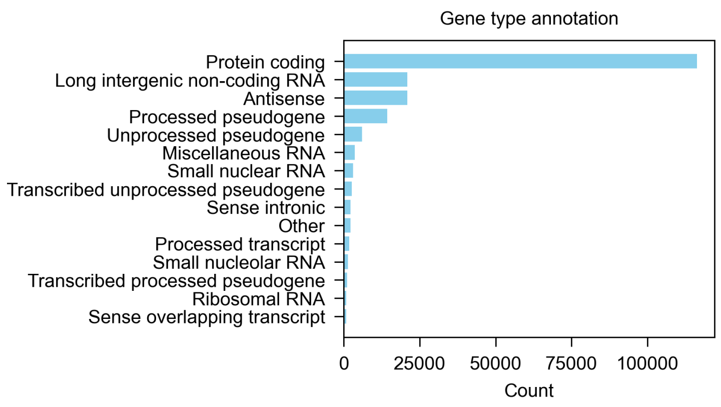


**Supplementary Fig. 4 | Gene type annotation of high sensitivity genes.** GENCODE v26 gene type annotation of genes with high gradients from all 20 models. Genes were sorted by mean saliency vector and the top 10,000 genes were selected from each model.

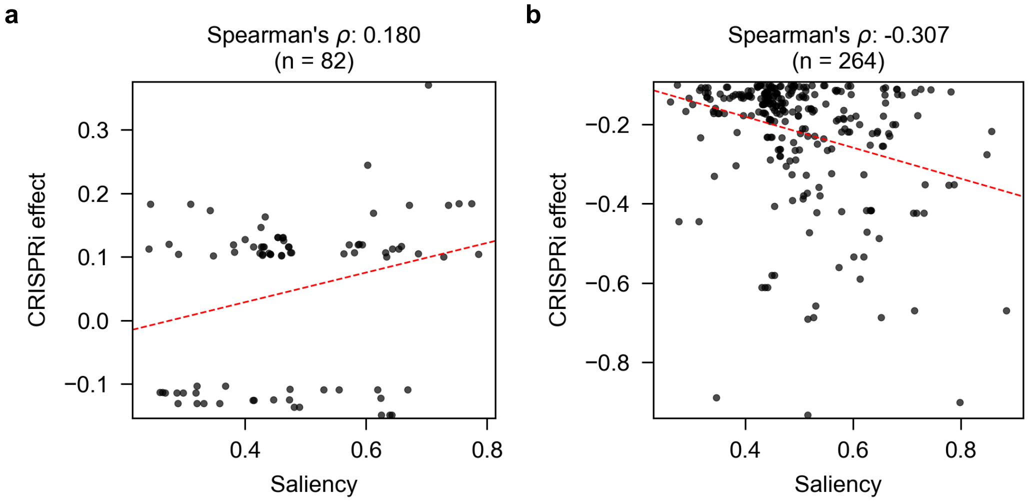


**Supplementary Fig. 5 | Sensitivity of experimentally validated CREs.** Correlation between summed saliency and CRISPRi predicted effect for high impact (absolute CRISPRi effect >10%) CRE-gene pairs. **a**, comparison between saliency and statistically insignificant cCREs. **b**, comparison between saliency and statistically significant cCREs. Negative correlation indicates better predictive power.

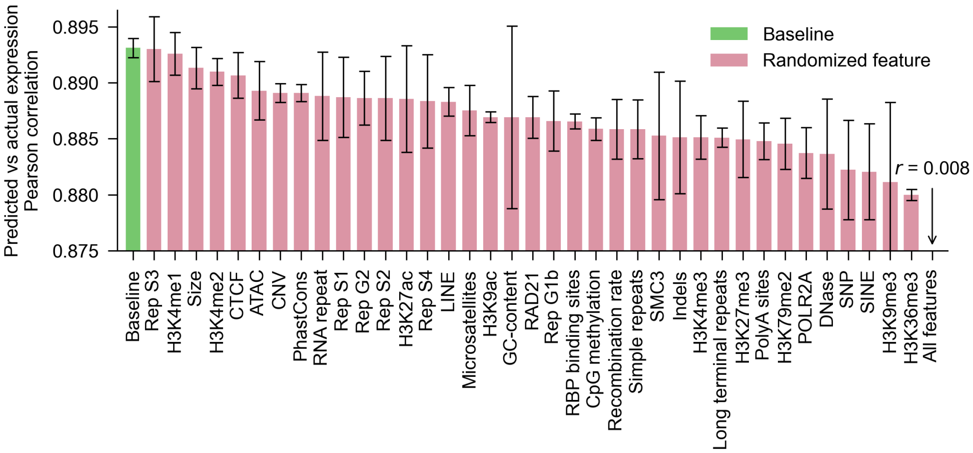


**Supplementary Fig. 6 | Predictive performance of models trained with randomized features.** Predictive performance of feature-randomized K562 models measured by averaged performance on hold-out test genes across three separate seeds. Node features of the specified type were randomized for the entirety of the graph. Error bars specify standard error across the three runs. The baseline model (green) utilizes all features without modification whereas other models (pink) randomize the specified feature. A model with all randomized node features was trained, but performance is below the visualization range (Pearson $R = 0.008$).

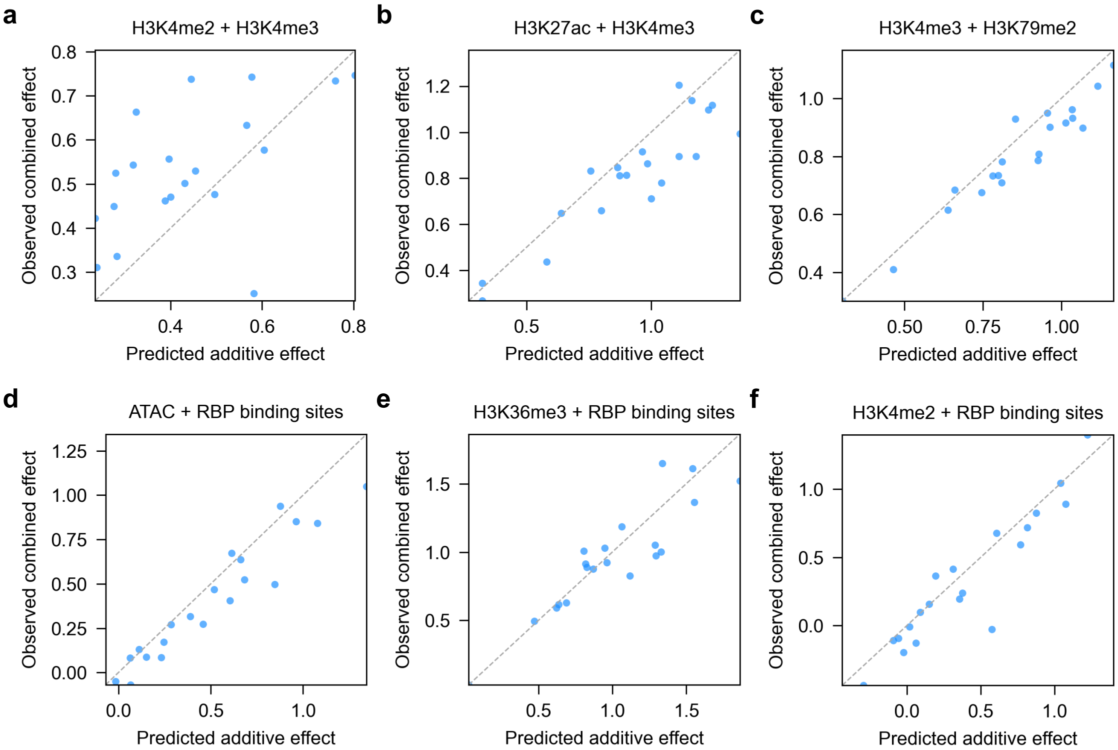


**Supplementary Fig. 7 | Joint feature ablations reveal non-additivity.** Lorem Ipsum

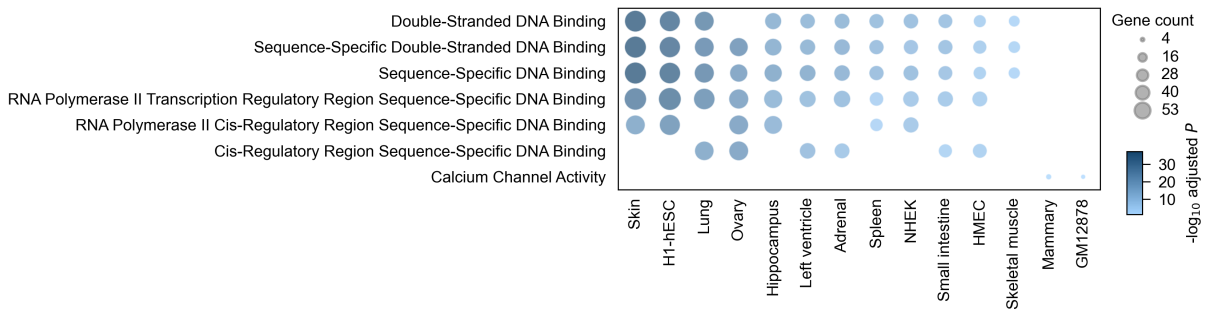


**Supplementary Fig. 8 | Gene set enrichment analyses of most affected genes from full H3K27me3 ablation.** Gene set enrichment analyses within the Go Molecular Function 2023 gene set. Analyses was run with the 100 most affected genes after full node feature ablation. Only gene sets with adjusted p-values < 0.05 and pathways in >= 2 tissues are shown.

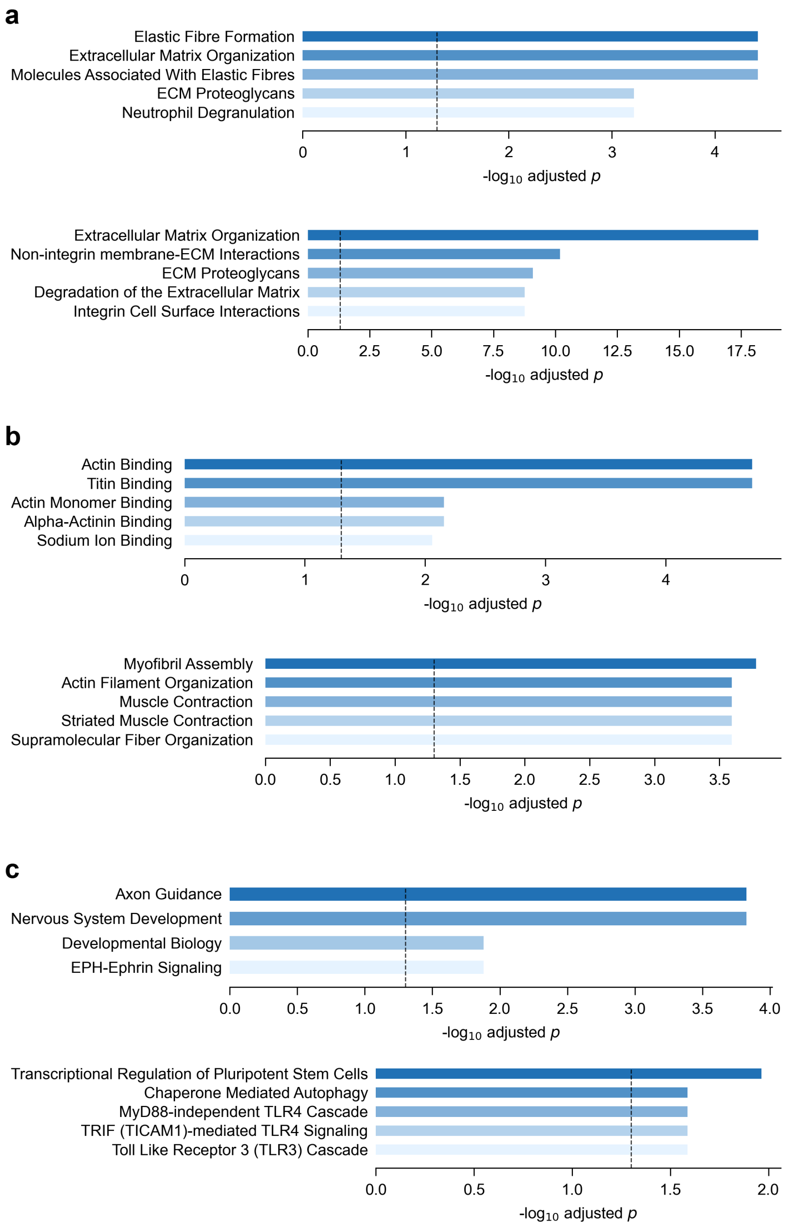


**Supplementary Fig. 9 | Sample-specific gene-set enrichment of most affected genes from full node feature ablation.** Each figure shows the results of running gene set enrichment analysis on the top 100 genes after full node feature ablation. Dashed lines indicate adjusted *p*-value = 0.05. **a**, Ablation of H3K27ac (top) and H3K36me3 (bottom) in the IMR90 model reveals an enrichment of genes involved with the extracellular matrix, **b,** Ablation of ATAC annotations (top) and H3K4me3 (bottom) in the skeletal muscle model reveals processes involved in motor unit control and organization. **c**, Ablation of DNase annotations (top) and H3K4me3 (bottom) in H1-hESC reveal processes involved in pluripotency and development. Gene sets used are **a**, **b**, Reactome Pathways 2024 and **c**, GO Molecular Function 2023 and GO Biological Process 2023.

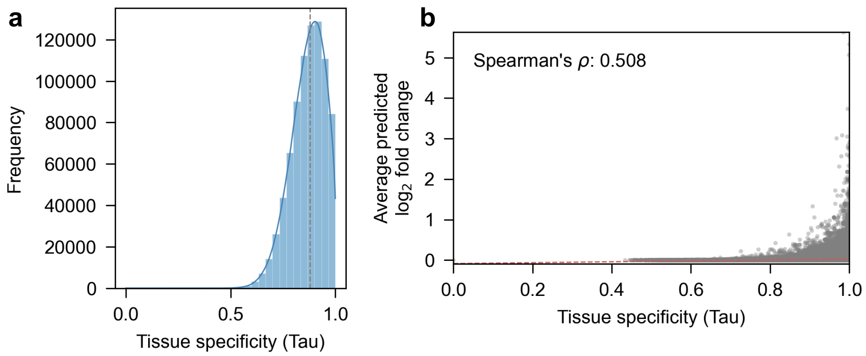


**Supplementary Fig. 10 | Tissue-specificity of node perturbation effects.** Histogram showing the distribution of tissue specificity (Tau) for perturbed node, **b**, Scatter plot of predicted relative fold change versus tissue specificity with Spearman's correlation coefficient $\rho = 0.508$.

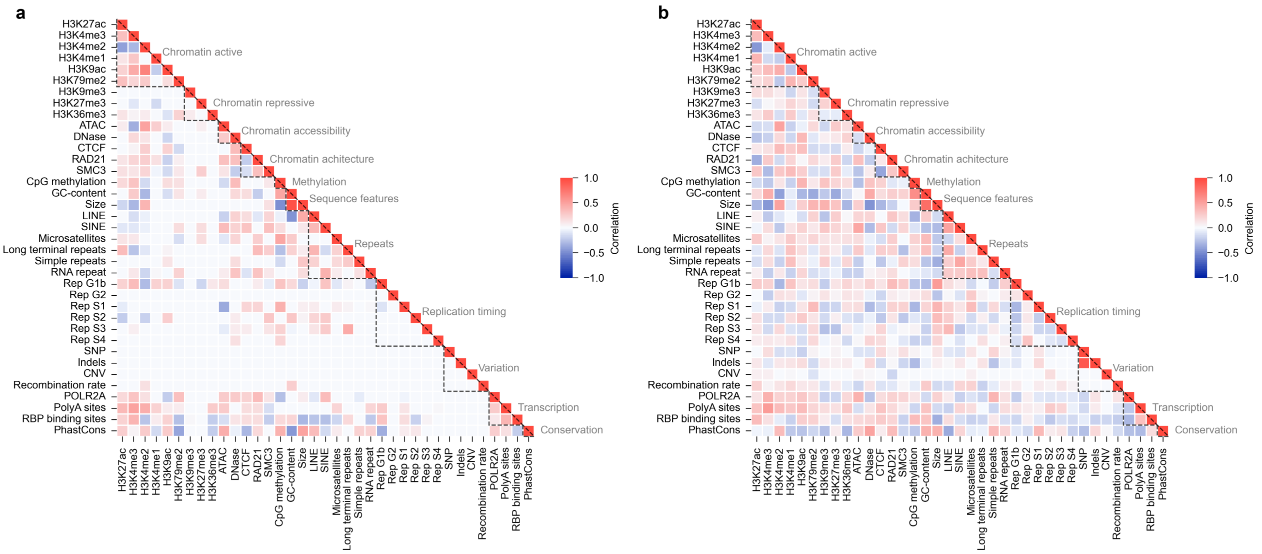


**Supplementary Fig. 11 | Partial correlations of feature effects reveal the cell-type differences in regulation.** Partial correlation matrices of feature ablation effects in a, left ventricle and b, HMEC, highlight cell-type specific patterns in feature utilization. While some relationships between features are conserved between cell types, many correlations are sample-specific, revealing the context-dependent nature of regulatory interactions. Features are organized by functional categories.

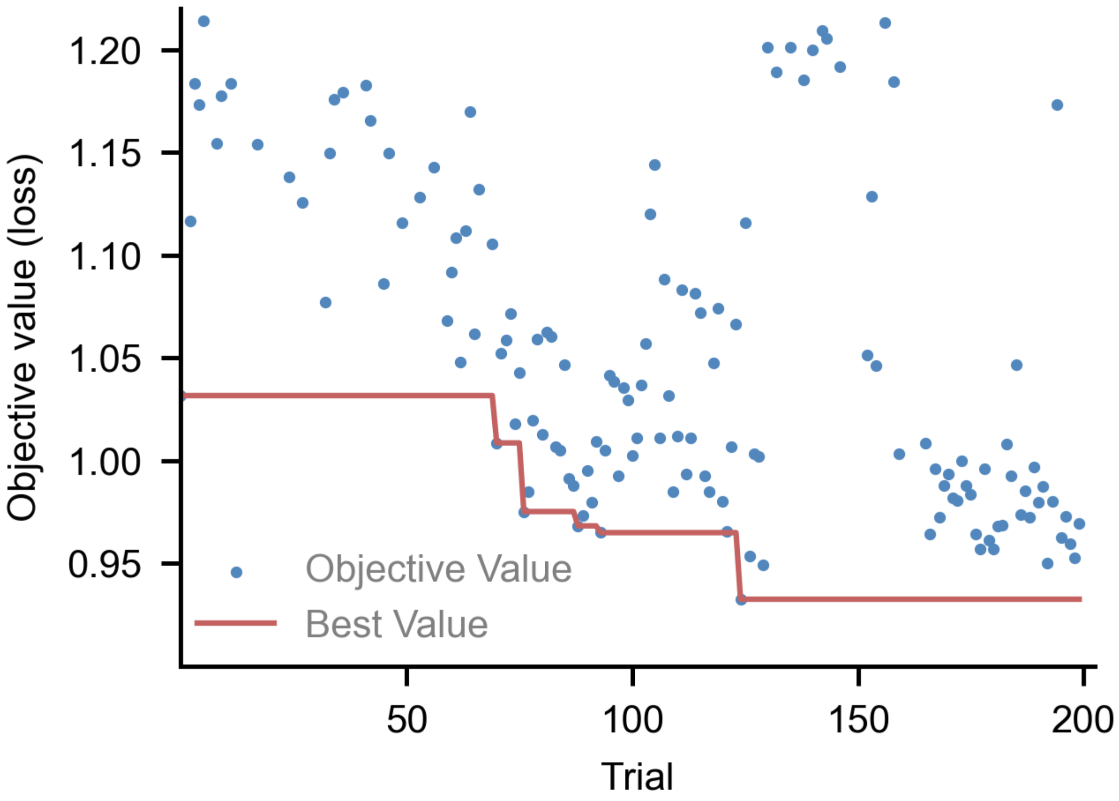


**Supplementary Fig. 12 | Optimization showing convergence over 200 trials.** Results of 200 optimization trials run on a subset of chromosomes. Results show validation RMSE decreasing with each pruned trial as params and architecture narrows adaptively to the graph construction specified. Blue dots indicate the results of individual trials, and the red line represents the minimum objective value found up to each point.

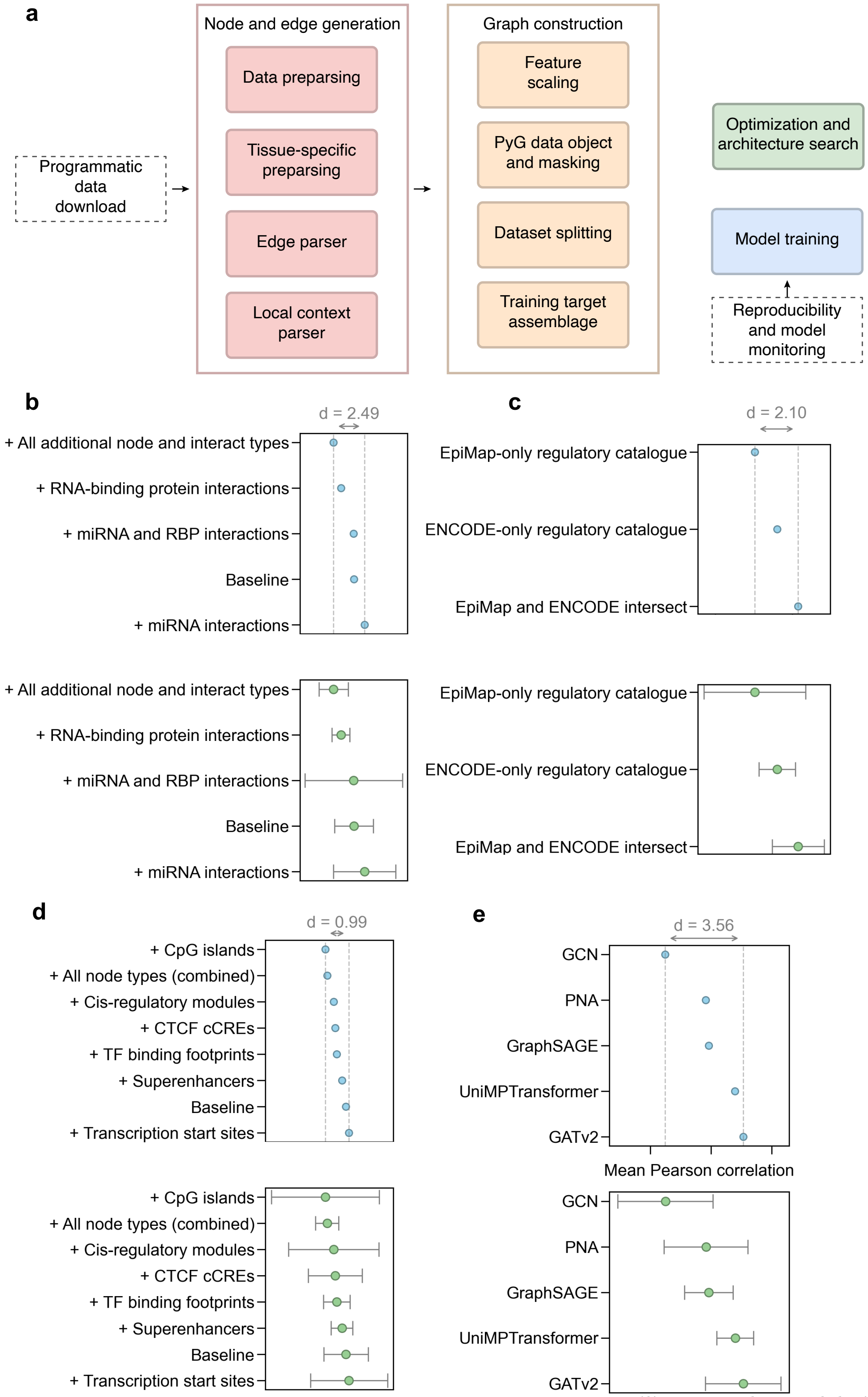


**Supplementary Fig. 13 | OGL's end-to-end design facilitates rapid iteration. a**, OGL utilizes an end-to-end design. After pre-processing, a single command takes the pipeline from data to completed optimization and NAS, or a trained model run over three random seeds for reproducibility. **b – e**, a series of iterative graph experiments run to converge on our most performant architecture. Plots on top show mean Pearson correlation across three random seeds, and bottom show mean bootstrapped Pearson (10,000 samples) across three random seeds on the validation set. **b**, adding different interaction data to the base graph, **c**, constructing graphs with varying regulatory element catalogues, **d**, adding different node types for graph construction, **e**, adjusting the GNN operator.
